# The potential of H5N1 viruses to adapt to bovine cells varies throughout evolution

**DOI:** 10.1101/2024.11.29.626120

**Authors:** Matthew L Turnbull, Mohammad Khalid Zakaria, Nicole S Upfold, Siddharth Bakshi, Callum Magill, Udeet Ranjan Das, Andrew T. Clarke, Laura Mojsiejczuk, Vanessa Herder, Kieran Dee, Nancy Liu, Monika Folwarczna, Georgios Ilia, Wilhelm Furnon, Verena Schultz, Hanting Chen, Ryan Devlin, Jack McCowan, Alex L Young, Wai-Wai Po, Katherine Smollett, Muhammad Ahsan Yaseen, Rebecca Ross, Avanti Bhide, Bianca van Kekem, Ron Fouchier, Ana da Silva Filipe, Munir Iqbal, Ed Roberts, Joseph Hughes, Dirk Werling, Pablo R. Murcia, Massimo Palmarini

## Abstract

Avian influenza H5N1 clade 2.3.4.4b viruses caused a global panzootic and, unexpectedly, widespread outbreaks in dairy cattle, therefore representing a pandemic threat. To inform effective control strategies, it is critical to determine whether the potential to adapt to bovine cells is a generalised feature of H5N1 viruses, or is specific to clade 2.3.4.4b, or even more restricted to specific genotypes within this clade (e.g., B3.13 and D1.1). Using a large panel of H5N1 viruses representing >60 years of their natural history and other IAV for comparative purposes, we demonstrate that virus adaptation to bovine cells is: (i) highly variable across 2.3.4.4b genotypes, (ii) limited in viruses predating the global expansion of this clade, (iii) determined by the viral internal gene cassette, and (iv) not restricted to udder epithelial cells. Mutations in the PB2 polymerase subunit, particularly M631L, emerge as key determinants of adaptation, although their phenotypic effects are context dependent and have limited enhanced viral polymerase activity in human cells. Bovine B3.13 and some avian genotypes also exhibit enhanced modulation of bovine interferon-induced antiviral responses, determined by at least the viral PB2, nucleoprotein, and the non-structural protein NS1. Our results highlight the polygenic nature of IAV host range and reveal that the potential to cross the species barrier varies during the evolutionary trajectory of H5N1, with some avian viruses more predisposed to spillover than others.

## INTRODUCTION

Influenza A viruses (IAV) represent a major threat to human and animal health. Wild aquatic birds are the main natural reservoir of IAV as they maintain multiple antigenically and genetically distinct subtypes of their external glycoproteins^1^ (17 hemagglutinins, HA, and 9 neuraminidases, NA). Humans, pigs, and horses instead are hosts to a limited number of IAV subtypes, originating from birds, which circulate in an endemic cycle within each species after pandemic/panzootic emergence (e.g. H1N1 and H3N2 in humans)^1^.

Certain avian IAV subtypes including H5Nx (where x represents distinct N subtypes), H7N9, H9N2, H10Nx and others, have occasionally infected some mammalian species including humans. These spillover events almost never lead to documented mammal-to-mammal transmission chains, but they nevertheless represent a potential pandemic risk, considering that each of the four influenza human pandemics that have occurred since 1918 were caused by viruses carrying at least some genomic segments from avian influenza IAV^2–6^.

Within this context, highly pathogenic H5N1 viruses are of particular concern as they are the cause of major devastating outbreaks in poultry, and have spilled over repeatedly in humans and other mammals^7^. Since 2021, H5N1 viruses within clade 2.3.4.4b have spread worldwide at unprecedented levels^8,9^, infecting a variety of mammalian species such as minks, skunks, raccoons, cats, bears, foxes^10–13^ and aquatic mammals including harbour seals and sea lions^14–17^. As these viruses spread, they reassorted with co-circulating low pathogenic avian IAV resulting in >80 genotypes so far, differing from each other by the constellation of their “internal” genomic cassette (i.e. the viral genome segments encoding all the viral proteins but HA and NA). Unexpectedly, in March 2024, H5N1 (B3.13 genotype) caused an influenza outbreak in dairy cattle in the US^18^. Until these outbreaks, cows had not been regarded as natural hosts of IAVs despite previous studies detecting anti-IAV antibodies in cattle sera, and experimental inoculations showing that cows are susceptible to both human and avian IAVs^19^ (reviewed in^20^). Affected cows showed clinical signs such as fever, depression, drop in milk production and mastitis, but only rarely signs of mild respiratory infections. Indeed, high levels of infectious virus could be detected consistently in the milk of infected animals^18^. The B3.13 genotype emerged after a European-origin 2.3.4.4b (EA-2020-C) entered North America in 2021^21,22^, and acquired PB2, PB1, NP and NS segments via sequential reassortment events with local low pathogenic avian influenza viruses (LPAIV)^22–25^. Phylogenetic analysis supports the notion that a single introduction event from wild birds into cattle occurred in late 2023 or early 2024 and this has subsequently spread to multiple dairy farms across 17 US states^26,27,28^. In January 2025, the emergence of a distinct genotype (D1.1) was reported in cattle. In the case of the D1.1 outbreak, a European-origin 2.3.4.4b virus had instead acquired the PB2, PA, NP and NA segments from local LPAIV via reassortment^24,29^. Bovine H5N1 spilled over to humans (70 confirmed human cases in the US as of August 2025^27^), in addition to cats and other small mammals in the vicinity of affected farms. Furthermore, a 2.3.4.4b virus of a different genotype (B3.6) caused a limited outbreak in goats^30^.

There are several known barriers to avian IAV adaptation in mammalian cells (reviewed in^31^), and cellular receptor specificity is a critical one. Avian HAs preferentially bind glycans that terminate with an α2,3-sialic acid, which are abundant in the gastroenteric tract of birds, while human-adapted IAV HAs preferentially bind to the α2,6-sialic acid conformation, which is abundant in the upper respiratory tract of humans and other mammals^32–36^. Therefore, a switch to efficient α2,6-sialylated glycan binding is considered an important prerequisite to successful avian-to-human adaptation. Consistent with this, several recent studies have focused on the bovine H5N1 virus glycoproteins and their receptor preference and tissue tropism^37–46^. α2,3-linked glycans, which avian HA can utilise, are abundant in the udder tissue of dairy cattle^38–40^, the primary site of infection with 2.3.4.4b bovine H5N1. The HA of 2.3.4.4b H5N1 virus associated with both bovine and human infections of the recent dairy cattle outbreak appears to retain a preference for ‘avian like’ α2,3-sialyl glycans^41–43^, although ability to bind α2,6-sialyl glycans to an extent has been reported^47^. Importantly, only a single amino acid change appears to allow the HA from an IAV isolated from a human dairy farm worker to bind α2,6-sialic receptors efficiently^44,45^, highlighting the potential for human-adaptation. Other glycoprotein-associated determinants hampering avian IAV transmission to mammals include pH sensitivity of HA, and the length of the NA glycoprotein stalk^48,49^.

Viral proteins encoded by the internal genomic segments (i.e. the PB2, PB1, PA, NP, M, and NS) also constitute key molecular determinants of virus host range. For example, polymerase complexes of avian IAV exhibit reduced efficiency in utilising mammalian ANP32 proteins, which are essential cofactors for IAV replication^50–55^, and the nuclear import machinery^56^. The nucleoprotein (NP) of avian viruses is the target of interferon stimulated genes MXA/ MX1 and BTN3A3, which human-adapted IAVs evolved to escape^57–62^. The non-structural protein 1 (NS1) and PA-X are multifunctional proteins, that counteract the IFN response and are also associated with host-range^57,63,64^.

In this study, we wanted to address whether adaptation to bovine cells was a generalised feature of H5N1 viruses, or if instead it was either a trait of viruses within the 2.3.4.4b clade or narrowly confined to specific genotypes within this clade. Given the zoonotic nature of H5N1 IAV, the size of the global cattle population (>1 billion^65^) and the high rate of contact between humans and cattle (and cattle-derived products), it is essential to identify the viral determinants that facilitate bovine adaptation of IAV. Here, we focused specifically on the internal gene cassette capturing over 60 years of evolution of avian IAV, as well as phylogenetically distinct viruses isolated from a variety of hosts and spillover events for comparative purposes. We tested a set of 89 reassortants carrying identical glycoproteins in a variety of *in vitro*, *ex-vivo* and *in vivo* assays to assess replication kinetics, polymerase activity, sensitivity to type-I IFN and restriction factors, and virulence in animal models.

## RESULTS

### Reassortants generated in this study

To investigate the molecular determinants of H5N1 adaptation to the bovine host, we first generated a set of 23 2:6 reassortants carrying the glycoproteins (HA and NA) of the laboratory-adapted H1N1 A/Puerto Rico/8/34 (PR8) strain, and the six internal genomic segments of various H5N1 viruses (Fig 1). Using reassortant viruses carrying identical glycoproteins allowed us to directly scrutinise the role of the viral proteins encoded by the internal gene cassette in adaptation to bovine cells.

**Fig 1.**
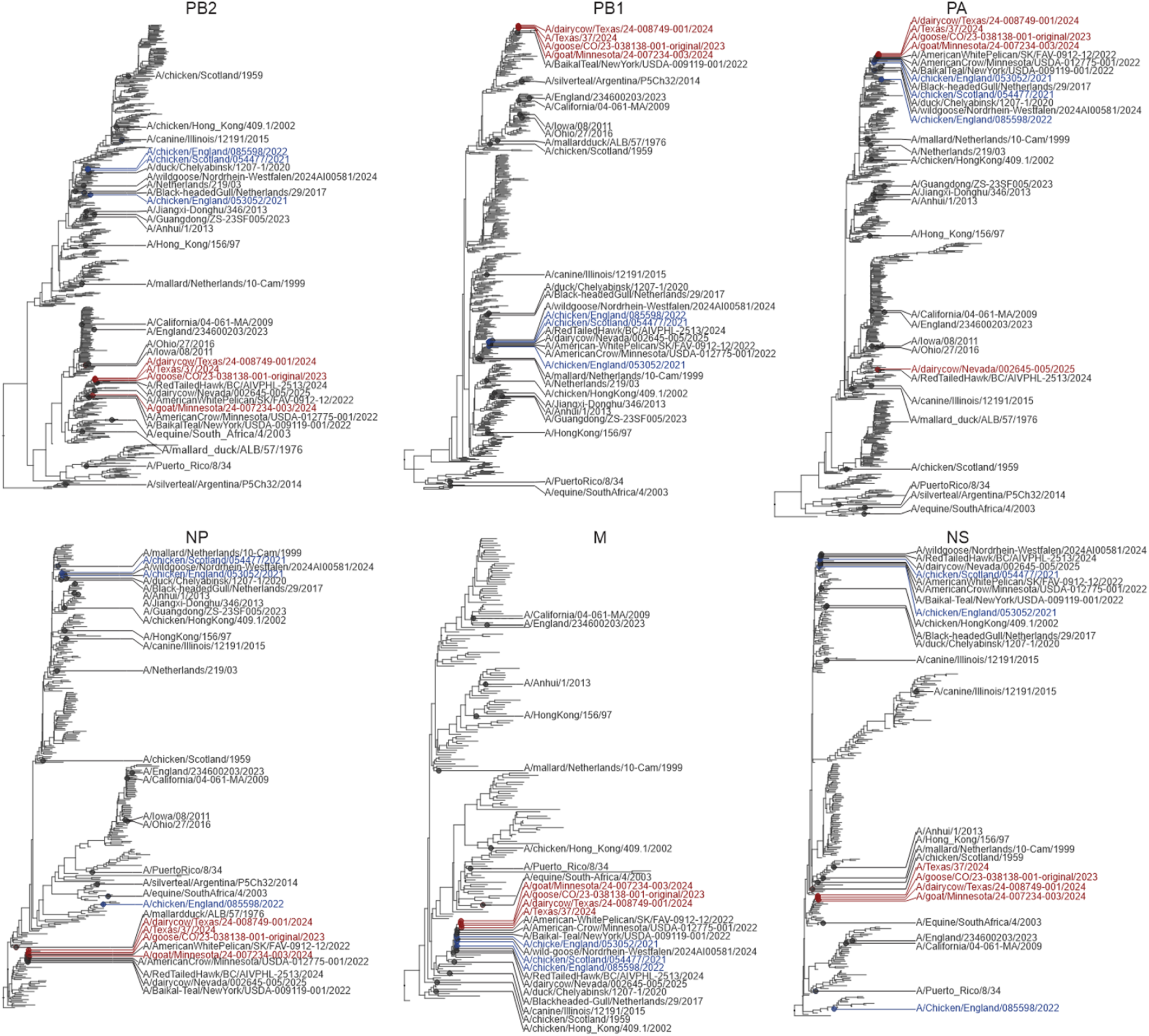
Global phylogenies of IAV sequences used in this study. Global maximum likelihood phylogenetic trees of nucleotide sequences from the internal segments of IAVs used in this study. Tips are annotated by strain names and correspond to viruses evaluated experimentally. Viruses belonging to the European and North American 2.3.4.4b clade are coloured blue and red, respectively.

Each reassortant used in this study is termed using the prefix “r” followed by an abbreviated name of the virus in question (Tables S1 and S2). For example, r-Bovine-B3.13 possesses the internal genes from one of the earliest sampled H5N1 bovine influenza sequences (sample acquired from a dairy cow on 20^th^ March 2024 in Texas). r-Texas/37-B3.13 is a similar, but distinct, B3.13 that infected a dairy farm worker and likely derives from a common ancestor of the lineage that has spread in cattle^26^. r-Bovine-D1.1 contains the internal genes of a distinct H5N1 outbreak of genotype D1.1 that was sampled in January 2025. r-Goat-B3.6 possesses the internal genes from the 2.3.4.4b virus isolated in goats (B3.6 genotype)^30^. We also generated reassortants with the internal genes of several other H5N1 2.3.4.4b genotypes isolated from birds from Europe and North America, including among those a reassortant representing the European virus that entered North America [r-EA-2020-C]. In addition, we also generated H5 viruses predating the global expansion of the H5N1 2.3.4.4b clade. These include the first sequenced HPAI H5N1 virus from 1959 [r-CHK/Scot/59]; a non-GsGd lineage of H5N2 from 1976 [r-Mld/76]; a clade 4 H5N1 from 2002 [r-CHK/02]; an early 2.3.4.4b H5N6 from 2017 [r-BHG/17]. For comparative purposes we included reassortants of the prototypic pandemic human 2009 virus [r-pdm09(H1N1)], and other mammalian adapted viruses (complete list of viruses used in Table S1).

We also generated 4:4 reassortants containing HA, NA, M and NS from PR8 and only the polymerase complex (PB2, PB1, PA and NP) from certain viruses of interest for specific assays (Table S2); these include some of the viruses indicated above, and others of avian [r-Arg(H4N2), r-Chn/23(H3N8), r-Mld/99(H1N1), r-Nld(H7N7), r-Chn/13(H10N8)], or swine [r-Iowa/11(H2N2v), r-Ohio/16(H3N2v)] origin. It should be noted that the virus from which the reverse genetics constructs were derived for Mld/99(H1N1) had been previously passaged in pig cells^66^. All reassortant viruses were rescued in HEK-293T cells, or in HEK-293T-Gg.ANP32A cells (293T cells expressing chicken ANP32) for those that could not be rescued in the parental cell-line (Table S1). Viral stocks were propagated and titrated in MDCK (or MDCK-Gg.ANP32A cells, Table S1) and fully sequenced as described in Methods.

### Viruses carrying internal genes derived from North American 2.3.4.4b ruminant viruses display the fastest replication kinetics in bovine cells

We first compared the replication kinetics of the panel of recombinant viruses in an IFN-competent cell line that we derived from bovine skin fibroblasts (BSF)^67^. r-Bovine-B3.13, r-Texas/37-B3.13, r-Goat-B3.6, and r-Bovine-D1.1 displayed the fastest replication kinetics, reaching higher titres at 24 hours post infection (hpi) than all other avian- and mammalian-origin IAVs tested. This included avian viruses ancestral to the B3.13 and D1.1 cattle outbreaks, as well as several distinct avian IAV genotypes. (Fig 2a-b & Supplemental Fig S1a-b). These “ruminant-derived” viruses reached high titres faster than other mammalian- and human-origin viruses, such as those isolated from equine or swine hosts, or following human spillover events. Importantly, the avian-origin recombinant viruses of the panel replicated as efficiently or better than r-Bovine-B3.13 in chicken DF-1 fibroblast cells (Fig 2c & Supplemental Fig S1c-d), and also relatively well in the permissive MDCK cell line (Fig S2a), suggesting this is a host species-specific phenotype and that intrinsic replication properties of the recombinant viruses (such as segment compatibility and RNA packaging) were not obviously attenuated. There seemed to be an inverse relationship between replication fitness in bovine and avian cells (i.e. ruminant viruses replicated well in bovine cells and less efficiently in avian cells and vice versa relative to the panel of avian-derived viruses). The only exception to this trend was r-Bovine-D1.1, which replicated to high levels in bovine and avian cells. Interestingly, we observed clear differences not only between the ruminant-origin viruses relative to the others, but also between different avian-origin IAVs. For example, viruses that replicated poorly in BSFs included reassortants derived from some 2.3.4.4b genotypes (e.g. EA-2020-C, B2.1, B3.1) and others that predate the origin of H5N1 2.3.4.4b viruses (i.e. r-CHK/Scot/59 (H5N1), r-Mld/76(H5N2), r-HK/CHK/02-Clade4 (H5N1) and r-BHG/17 (H5N6)) (Fig 2b&d). This overall trend was also observed in primary nasal fibroblasts (BNF) isolated from bovine nasal tissue received from an abattoir (Fig S2b), suggesting this was not a phenotype specific to skin fibroblast cells. Importantly, the replication advantage observed for r-Bovine-B3.13, a representative ruminant-origin IAV, was maintained in *ex vivo* bovine udder explant tissue from multiple donors. In contrast, avian-derived recombinant viruses showed reduced replication, supporting the reliability of the phenotype observed in the bovine skin fibroblast model (Fig 2e & f). This trend was also consistent across a range of multiplicities of infection when compared to the earlier European r-EA-2020-C virus which entered North America (Fig S3a). Of note, our 2:6 viruses appeared to replicate preferentially in epithelial cells of the cistern of the mammary glands (Fig 2g & Fig S3 b&c) as assessed by immunohistochemistry of fixed tissues. Although trends were clear, data was quite variable, and we carried out experiments using several replicates (18 replicates in total from tissue derived from 6 different donors) in order to produce statistically robust data. The variability of the data is determined by the technical difficulty to obtain explants with a homogenous size of cisternal epithelium from these explants.

**Fig 2.**
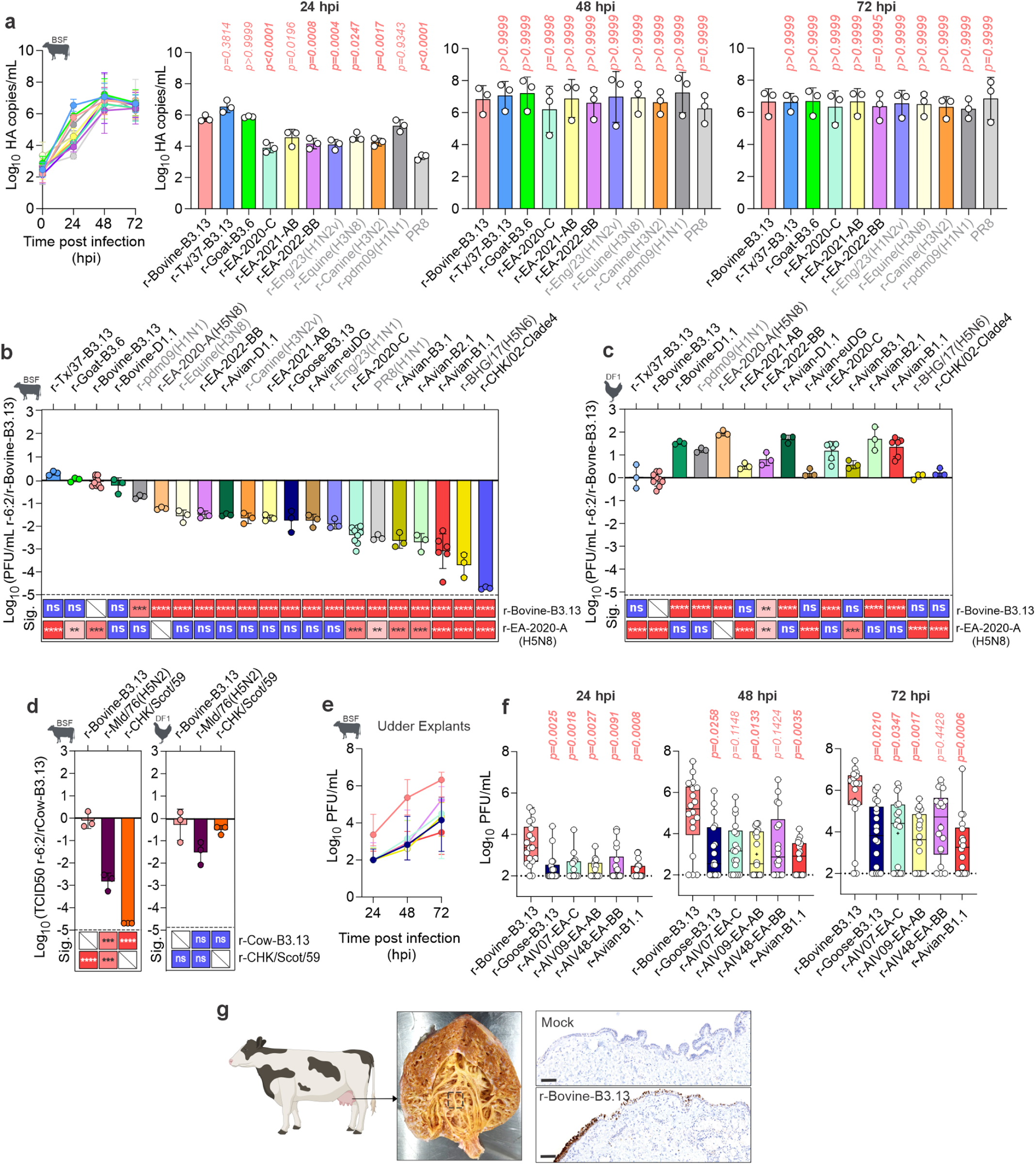
Replication of 2.3.4.4b and other IAV reassortant viruses in bovine cells and udder tissue. **a**, Bovine skin fibroblasts (BSF) were infected with 0.0005 viral genome copies per cell and copies in the supernatant at each time point were quantified by RT-qPCR. **b-d**, As above, but BSF (**b**) and chicken embryonic fibroblasts (DF1) (**c**) were infected with 0.001 PFU per cell and infectious virus in the supernatant at 24 hours post infection (hpi) was quantified by plaque assay on MDCK cells (**b** and **c**) or by TCID50 on DF1 cells (**d**). **e**-**f**, Bovine udder tissue explants were infected with 1000 PFU per explant. Virus released into the supernatant was quantified by plaque assay on MDCK cells. **g**, Photomicrographs of r-Bovine-B3.13 or mock-infected explant tissue derived from bovine udders. Cells positive for IAV nucleoprotein (N; shown in brown) are found in the epithelial layer of the tissue. In **a**-**d**, data are mean +/- s.d. of three independent biological experiments (n = 3). For (**a**) individual points represent the mean of two technical duplicates per biological repeat. In **a-d**, raw data (**a**) or data normalized to r-Bovine-B3.13 (**b-d**) were log-transformed, and normality was confirmed using the Shapiro-Wilk Test. At each timepoint, multiple comparisons between all viruses were performed using an ordinary one-way ANOVA with Tukey’s multiple comparison test (α= 0.05). Comparisons are shown relative to r-Bovine-B3.13(H5N1) in all panels, with additional comparisons to r-EA-2020-A (H5N8) or r-CHK/Scot/59 shown in **b-c** and **d**, respectively. Data in **e-f** are from three independently infected tissue explants from 6 biological donors (n = 18). These data were log-transformed but did not meet the assumption of normality (Shapiro–Wilk test). The median and interquartile range per virus and timepoint are shown in **e**, while individual explant titers per virus group are overlaid box-and-whiskers plots for each timepoint in **f**. Boxes represent the interquartile range, the line indicates the median, and whiskers show the full data range. Statistical significance between the r-Bovine-B3.13 virus and other virus groups were determined using the Kruskal–Wallis test followed by Dunn’s multiple comparisons. In all panels P-values in bold indicate statistical significance. In **b-d** ns = p > 0.05; * = p ≤ 0.05; ** = p ≤ 0.01; *** = p ≤ 0.001; **** = p ≤ 0.0001. H5N1 viruses are shown in black, while labels for non-H5N1 IAVs are shown in grey.

Overall, these data suggest that the internal genomic segments of viruses associated with recent outbreaks in ruminants in North America provide a replication advantage in bovine cells relative to other diverse avian- and mammalian-origin IAVs. As these viruses contain internal genes derived from three distinct genotypes (B3.6, B3.13 and D1.1), this suggests that more than one constellation of internal genomic segments can confer enhanced replication fitness in a bovine host. Furthermore, these data support the notion that replication potential of H5N1 viruses in bovine cells varies widely between lineages, clades and genotypes and it is independent from the udder microenvironment.

### “Bovine-specificity” of B3.13, D1.1 and B3.6 viruses does not directly correlate to high polymerase activity in human cells

The activity of the viral polymerase complex (PB1, PB2, PA and NP) is a critical determinant of IAV within-host fitness and therefore central for virus tropism and adaptation. As the viruses we assessed exhibited different replication kinetics, we performed standard minireplicon assays in human 293T cells to evaluate the specific contribution of the polymerase to the observed phenotype in mammalian cells. We tested the polymerase complexes of all the viruses of the 2:6 panel described above, as well as a variety of other control viruses (outlined in Tables S1, S2 and S3), to obtain a comprehensive and comparative overview of their activity in human cells (Fig 3a).

**Fig 3.**
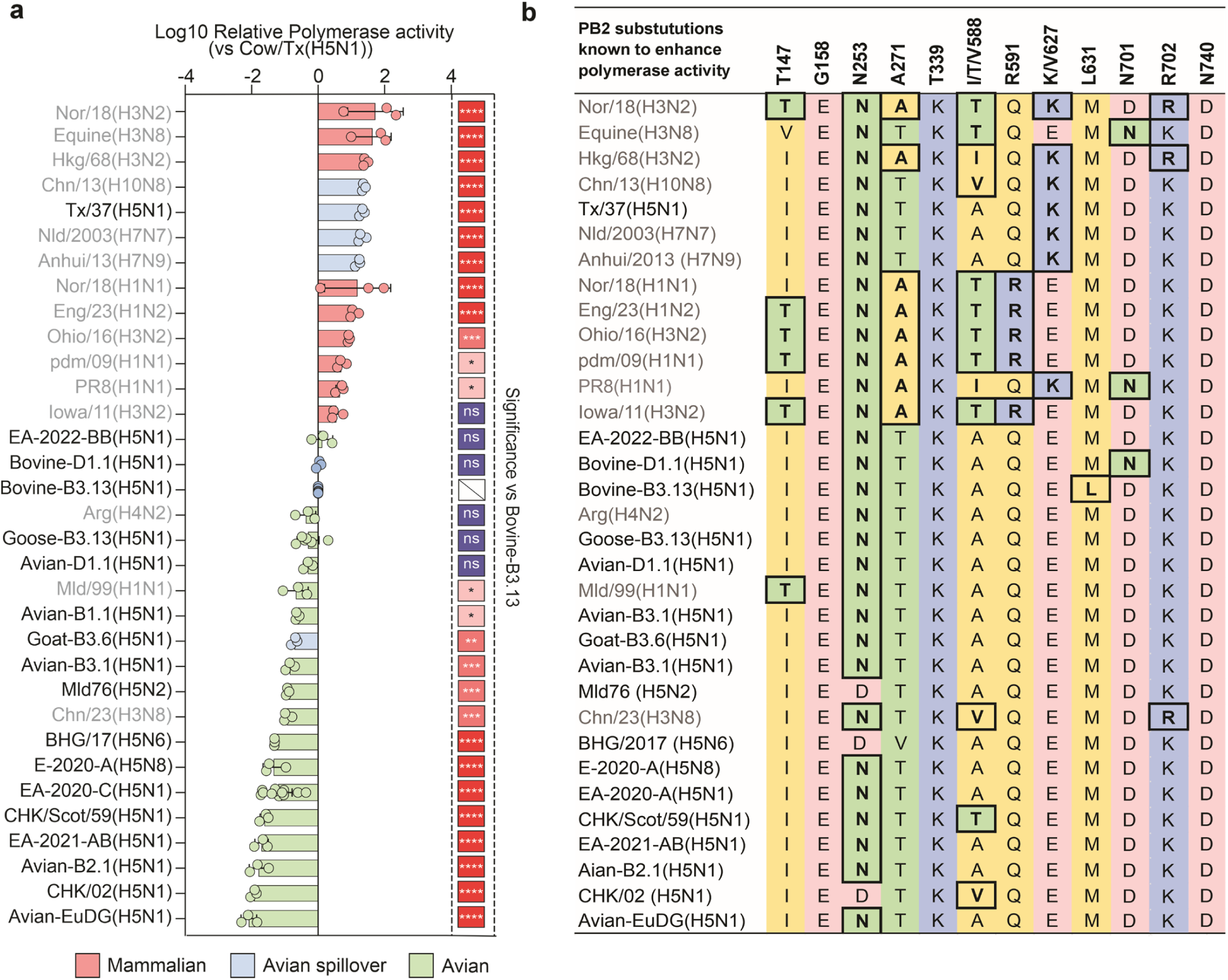
Polymerase activity of diverse IAV strains in human cells. Minireplicon assays were performed in HEK293T at 24 hours post transfection to assess the polymerase activity of a broad panel of IAV viruses. Data are mean +/- s.d. of at least three independent experiments and each data point represents the mean of two technical duplicates from each biological repeat. All data were normalized to r-Bovine-B3.13, log-transformed, and normality was confirmed using the Shapiro-Wilk Test. Multiple comparisons between all polymerases were performed using an ordinary one-way ANOVA with Dunnett’s multiple comparison test (α= 0.05). Comparisons are shown relative to r-Bovine-B3.13. Statistical symbols: ns = p > 0.05; * = p ≤ 0.05; ** = p ≤ 0.01; *** = p ≤ 0.001; **** = p ≤ 0.0001. H5N1 virus polymerases are shown in black, while labels for non-H5N1 IAVs are shown in grey. **b**, Table of PB2 residues that have been implicated in increased polymerase activity in previous studies. Black boxes encircle residues that enhance polymerase activity described in other studies^50,68–70,107,125–130^

The highest activity was reached with the polymerases from IAVs of human-origin or those associated with human spillover events. Among the top 13 ranked polymerases, all but an equine-origin virus and the laboratory-adapted PR8 were derived from human-endemic or spillover viruses of avian- or swine-origin. These viruses possess mammalian-adaptative mutations in PB2 including the well-described 627K, 701N and 591R substitutions, among others (Fig 3b)^50,68–70^. These viruses possess mammalian-adaptative mutations in PB2 including the well-described 627K, 701N and 591R substitutions, among others (Fig 3b)^50,68–70^.

Bovine-derived polymerases displayed intermediate activity (Fig 3a), while most avian-derived polymerases exhibited, as expected, the lowest recorded activity in mammalian cells. The sequence of PB2 derived from r-Bovine-B3.13 H5N1 does not possess any major known PB2 adaptive mutations, with the exception of M631L which has been reported to boost polymerase activity of a mouse-adapted H10N7 virus and has recently been shown to contribute to enhanced activity of a human-derived B3.13 polymerase in human cells^7,71,72^.

Some avian IAV polymerases, including the one derived from EA-2022-BB, which ranked highest among avian-origin polymerases, exhibited polymerase activity similar to those of bovine-adapted viruses. Notably, the EA-2022-BB genotype has been implicated in outbreaks among farmed fur mammals in Europe^73^. Other relatively high-ranking avian polymerases included Avian-D1.1 and Goose-B3.13, both closely related to viruses that emerged in cattle. The goat-derived polymerase displayed lower activity than other avian-origin polymerases, including Avian-D1.1 and Goose-B3.13.

Taken together, these data highlight that adaptation of certain H5N1 viruses to bovine cells is also correlated with a relative enhancement of polymerase activity to human cells, although this does not reach the levels of human-adapted viruses. Importantly, polymerases of some specific avian viruses also possess activity in human cells similar to bovine-adapted viruses. The data also suggest that polymerase activity alone is not necessarily the only predictor of virus fitness in replication assays, and that additional genomic segments may compensate for suboptimal polymerase activity.

### r-Bovine-B3.13 is more virulent than ancestral r-EA-2020-C in mice

We next inoculated C57BL/6 mice with 100 PFU of either r-Bovine-B3.13, r-EA-2020-C or PR8 as control, to test whether the internal genes of bovine B3.13 virus were sufficient to drive viral spread in respiratory tissues and determine virulence in this widely used experimental model for IAV^74^. As expected, PR8 (a mouse-adapted laboratory strain) caused severe weight loss in infected animals, while the weight of mice infected with r-EA-2020-C was essentially unaffected. Mice infected with r-Bovine-B3.13 displayed an intermediate phenotype showing weight loss but not as severe as those infected with PR8 (Fig S4a). Both quantification of NP in the lungs of infected mice euthanised at 3 days post-infection (dpi) and T-cell infiltrates (CD3) in mice euthanised at 8 dpi displayed similar trends, with higher values in mice infected with PR8, followed by those infected with r-Bovine-B3.13 and then r-EA-2020-C, although differences were not statistically significant due to variability within groups (Fig S4b-d). As highly pathogenic H5N1 can induce a systemic infection, we assessed the presence of NP in the brains of infected mice to determine virus spread in organs other than the lungs. We could not detect viral NP by immunohistochemistry in any brain of the infected animals (not shown). Lack of systemic infection by r-Bovine-B3.13 was to be expected as HPAI HA is a key determinant of systemic infection^75,76^. Thus, the internal genes of bovine B3.13, unlike those of the avian EA-2020-C genotype, are sufficient to induce some pathology in mice, although not as pronounced as those induced by wild type B3.13 in published studies^77,78^.

### Synergistic interactions among internal genomic segments of B3.13 viruses contribute to their higher replication fitness in bovine cells

Our data suggested that polymerase activity measured in minireplicon assays is not a stand-alone predictor of replicative fitness of live viruses in bovine cells. To directly assess the role of the polymerase complex during viral replication, we infected bovine (BSFs) and avian (DF1) cells with PR8-recombinant viruses harbouring the polymerase and NP genes from select viruses (referred to as 4:4 viruses).

When 4:4 viruses were compared in BSF cells, the replicative fitness of the viruses in the panel was different from their fitness as 2:6 viruses (which harboured their cognate M and NS segments, Fig 4a-c). Whilst r-Bovine-B3.13 was amongst the fittest in this panel of 4:4 viruses in BSF cells, there was no longer a clear advantage over the avian r-EA-2022-BB 4:4 virus, as observed in the context of the 2:6 assays (Figs 2 a & b), which in turn displayed similar polymerase activity to the bovine B3.13 polymerase in minireplicon assays (Fig 3A). Furthermore, the r-Goat-B3.6 4:4 virus, which displayed one of the fastest replication kinetics as a 2:6 virus (Figs 2 a & b), replicated at similar levels to those observed by avian-origin viruses r-EA-2020-C and r-Goose-B3.13. Importantly, all the 4:4 viruses replicated as well as, or in the case of r-EA-2022-BB better than, the r-Bovine-B3.13 4:4 virus in chicken DF-1 cells. Overall, while these data are line with the results obtained in human polymerase assays, they suggest also that M and/or NS play an important role in virus replication in bovine cells.

**Fig 4.**
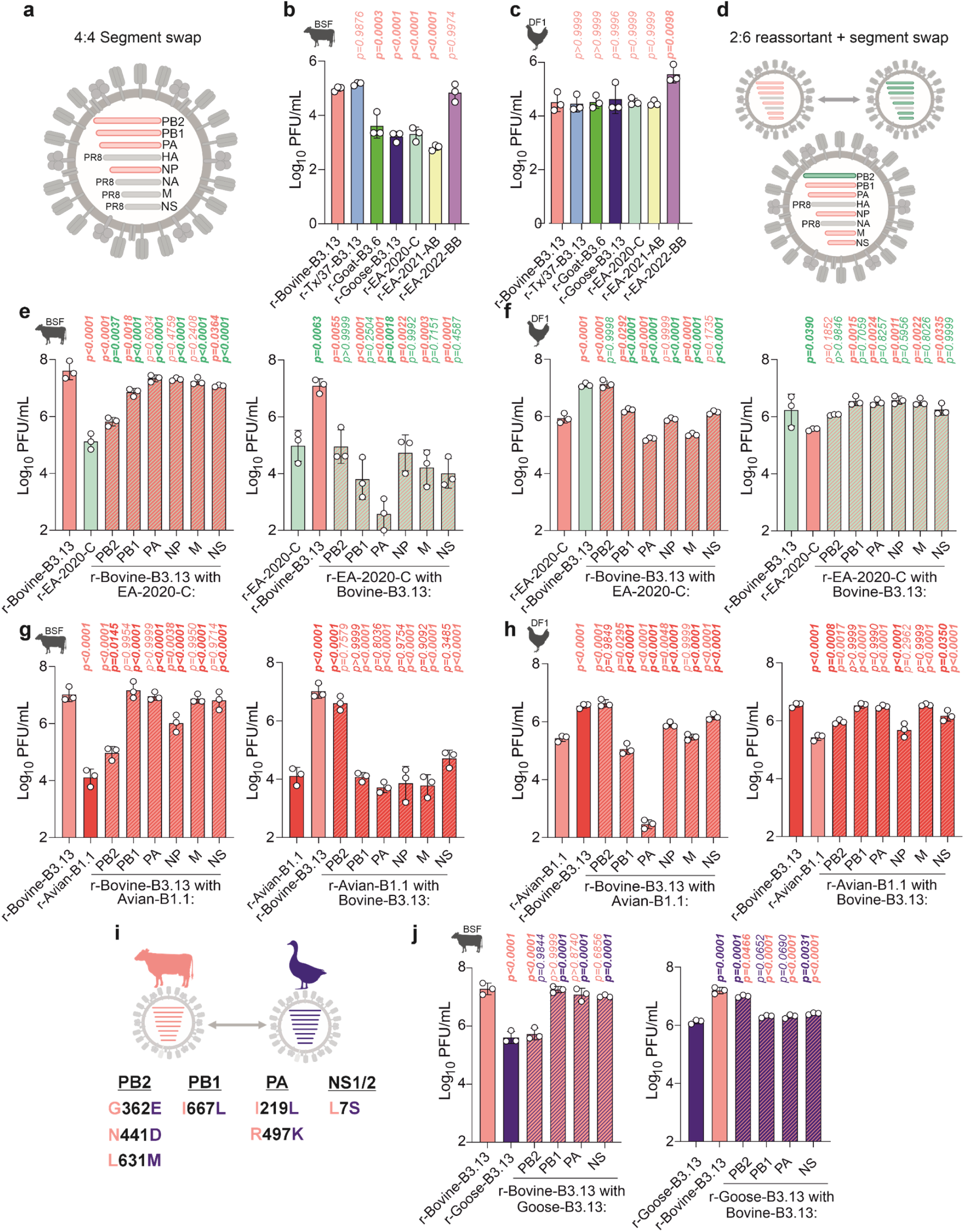
Segment-specific contribution of B3.13 internal genes to virus replication. **a**, Schematic of a 4:4 reassortant virus containing segments 1,2,3 and 5 from an H5N1 virus in the PR8 backbone. **b-c**, Replication of 4:4 recombinant viruses following infection of BSF (**b**) or DF1 (**c**) for 24 hpi. **d**, Schematic of 2:6 viruses used in **e-h**, with internal gene segments reciprocally exchanged between r-Bovine-B3.13 and either r-EA-2020-C or r-Avian-B1.1. **e-h**, Replication of 2:6 reciprocal viruses in BSF (**e,g**) and DF1 (**f,h**) cells at 24 hpi. **i**, Schematic of amino acid changes between r-Bovine-B3.13 and r-Goose-B3.13. **j**, Replication of 2:6 reciprocal viruses between r-Bovine-B3.13 and r-Goose-B3.13 in BSF cells. An MOI of 0.001 PFU/cell was used in all infections, and subsequent virus released into the supernatant at 24 hpi was titrated by plaque assay on MDCK cells. Data are mean +/- s.d. of three biological repeats. Data were log-transformed and confirmed to be normally distributed using the Shapiro-Wilk Test. Multiple comparisons between all viruses were performed using an ordinary one-way ANOVA with Tukey’s multiple comparison test (α = 0.05). Only comparisons relative to controls are shown. P-values in bold indicate statistical significance.

To dissect further the role(s) of individual segments, we first made individual reassortants between r-Bovine-B3.13 and r-EA-2020-C (Fig 4d), representing the European genotype that first entered North America. There are multiple changes across the genome between these two viruses (see Supplementary File 1 for an alignment of all internal gene proteins from avian H5Nx viruses used in this study relative to r-Bovine-B3.13). The NP protein has the fewest of the non-spliced products (3 substitutions) while each of the polymerase subunits has at least 10 changes each. When individual segments of the bovine virus were replaced with those from avian r-EA-2020-C, there was a significant reduction in titre for the PB2, PB1 and NS segment reassortants, with PB2 causing the largest decrease relative to the bovine control (Fig 4e). None of these individual swaps reduced the titre to the level of the r-EA-2020-C control, however. In the reciprocal reassortant viruses, none of the internal segments from the bovine B3.13 reassortant increased replication of r-EA-2020-C virus in bovine cells. Indeed, only when four segments of bovine B3.13 (PB2, PB1, NP and NS) were introduced together into EA-2020-C, reflecting the four segments derived from North American LPAI during the emergence of B3.13, there was a clear effect on the replication of the EA-2020-C parent (Fig S5). Importantly, reassortant viruses that were attenuated relative to the r-Bovine-B3.13 control in bovine cells, replicated to high titres in chicken cells, suggesting these viruses did not have an obvious intrinsic defect (such as genome packaging with the PR8 HA and NA segments) (Fig 4f).

We next rescued a set of reassortants between r-Bovine-B3.13 and r-Avian-B1.1, a virus that replicated particularly poorly in BSF cells and udder tissue (Fig 2b) and is a distinct American genotype acquiring PB2, PB1 and NP from local LPAIV^79^. As with EA-2020-C, there were multiple changes across the genome relative to r-Bovine-B3.13 (Supplemental File 1) with PB2 having the greatest number of substitutions (13) and NP the fewest (4) of the non-spliced genes. In these reassortants, the avian B1.1 PB2 reduced the replication of r-Bovine-B3.13 significantly, although not the level of the r-Avian-B1.1 control, but did not reduce titres in chicken DF-1 cells (Fig 4g & h). The avian B1.1 NP segment also modestly reduced the replication of r-Bovine-B3.13. In the reciprocal setup, bovine PB2 rescued r-Avian-B1.1 replication to levels similar to the r-Bovine-B3.13 control, suggesting a more dominant role for PB2 in this virus context.

Finally, we explored the contribution of individual segments to the phenotypic differences observed between bovine B3.13 and the reassortant derived from the closest sequenced B3.13 virus isolated from birds (r-Goose-B3.13). As there are only a small number of amino acid changes in PB2, PB1, PA and NS (Fig 4i), only these segments were swapped between viruses. In bovine cells, recombinant viruses with either goose-B3.13 PB1, PA or NS in r-Bovine-B3.13 reached titres similar to r-Bovine-B3.13. Importantly, the r-Bovine-B3.13 reassortant bearing the PB2 segment from goose-B3.13 fully recapitulated the replication phenotype of r-Goose-B3.13 (Fig. 4j). These data suggest that PB2 adaptive mutations played a particularly important role in improving virus replication after emergence in dairy cattle, in accordance with previous studies and observations^72,80^.

Overall, our segment swap data highlight that efficient replication in bovine cells requires multiple adaptive mutations, likely due to the presence of multiple, distinct bovine barriers to avian influenza viruses. The number, position, and nature of adaptive mutations are likely to vary depending on the genomic constellation of the virus genotype.

### PB2 M631L is an important adaptive mutation for bovine B3.13 virus in bovine cells but is context dependent

Our data suggested that adaptive mutations in PB2 were important in defining phenotypic differences before- and after-emergence of B3.13 in cattle from an avian host. Importantly, only three amino acid residues differentiate the PB2 proteins of r-Goose-B3.13 and r-Bovine-B3.13 (E362G, D441N, M631L; Fig 4i and Fig 5a). Of those, the mutation PB2 M631L was previously shown to be an adaptive mutation in mammals^71,72^. Indeed, only a single 631M mutation reduced the replication of r-Bovine-B3.13 to levels similar to r-Goose-B3.13 in BSF cells (Fig 5b). Introducing the PB2 M631L substitution did significantly enhance the replication of r-Goose-B3.13. However, it was not sufficient to fully recapitulate the phenotype conferred by the entire bovine PB2 as seen in Fig 4j or enhance replication to levels observed for r-Bovine-B3.13 (Fig 5b), suggesting that one of both of the additional substitutions, 362G and 441N, may further contribute to enhanced replication in bovine cells. Indeed, replication was similar to r-Bovine-B3.13 when either the E362G or D441N substitutions were individually introduced alongside M631L in the context of the Goose-B3.13 PB2 segment (Fig 5b). Interestingly, the PB2 D441L+M631L substitution combination resulted in the greatest enhancement of replication, reaching titres within a 2-fold difference of the r-bovine-B3.13 virus, although this was not significantly different to the E362G+M631L combination. In polymerase assays (Fig 5c), PB2 M631L alone boosted activity above that of the r-Bovine-B3.13 and r-Goose-B3.13 polymerases. Interestingly, the E362G+M631L combination boosted the r-Goose-B3.13 polymerase, but to levels below that of the single substitution. Conversely, the combination of D441N and M631L enhanced polymerase activity to levels far higher than that of the single substitution or parental polymerases. Together these results suggest that M631L plays an important role in bovine-adaptation, consistent with previous studies^72,80^, and further reveal that E362G and D441N provide an additive effect, enhancing replication beyond that of M631L alone in bovine cells.

**Fig 5.**
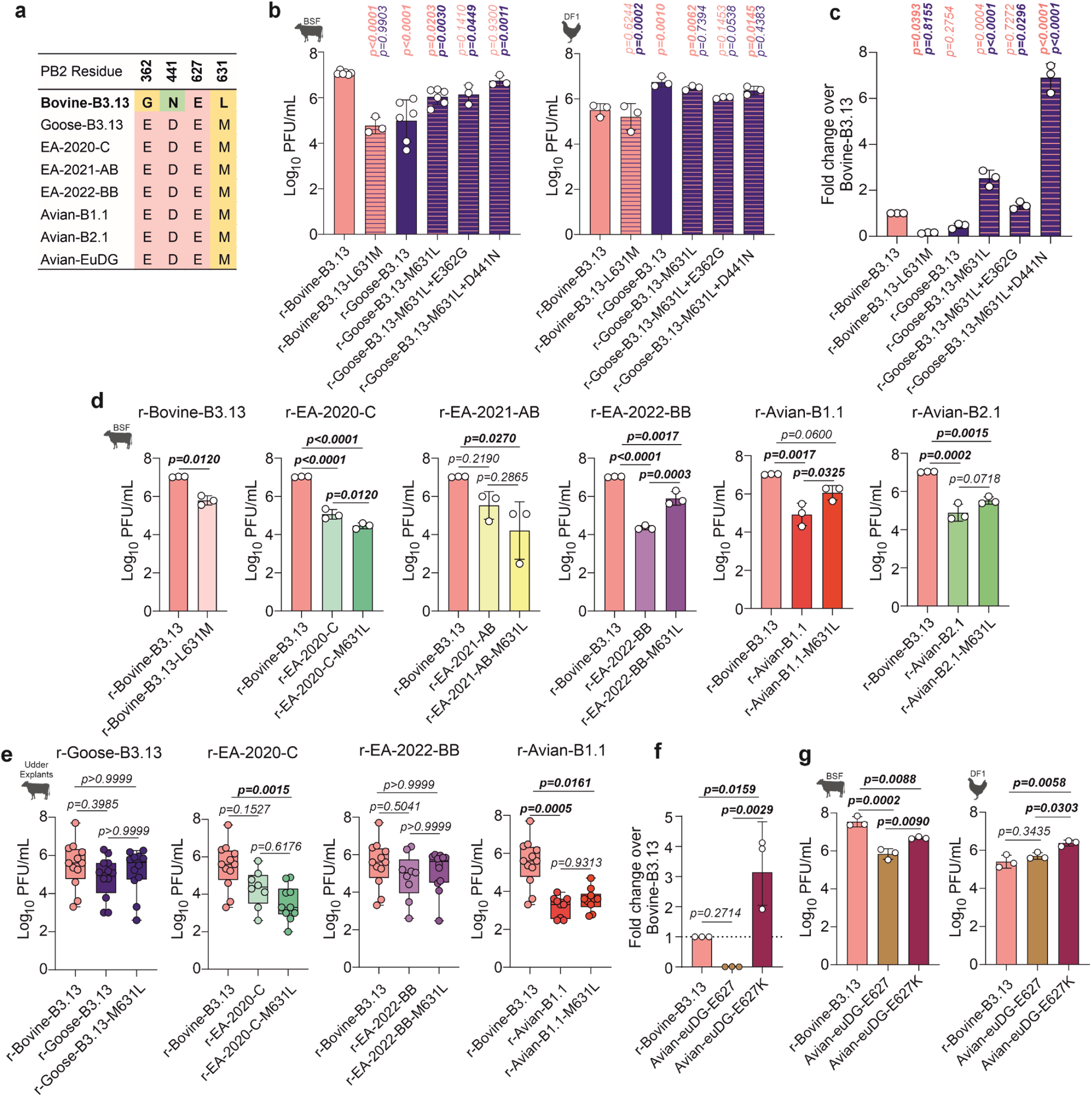
Replication of avian 2:6 viruses bearing M631L and other r-Bovine-B3.13 associated PB2 substitutions. **a**, Alignment of PB2 residues against r-Bovine-B3.13 (bold). **b**, BSF or DF1 cells were infected with r-Bovine-B3.13 or r-Goose-B3.13 WT or mutant viruses carrying individual or double PB2 amino acid substitutions from the reciprocal virus. Infections were performed at an MOI of 0.001 PFU/cell and virus in the supernatant at 24 hpi was titrated by plaque assay in MDCK cells. **c,** Minireplicon assays were performed in HEK293T at 24h post-transfection to assess the effects of PB2 amino acid substitutions in **b** on polymerase activity. Data are normalized to r-Bovine-B3.13 activity. **d-e**, Replication of WT or mutant r-Bovine-B3.13 and avian genotype representatives carrying the PB2-M631L substitution in BSF cells (**d**), or udder tissue explants (**e**). **f**, Minireplicon assays (as in **c**), comparing polymerase activity of r-Bovine-B3.13 and WT or PB2-E627K-bearing r-Avian-EuDG. **g**, Replication of r-Bovine-B3.13 and WT or PB2-E627K-bearing r-Avian-EuDG in BSF and DF1 cells at 24 hours post infection (as in **b**). In **b, d** and **f-g**, data are mean +/- s.d. of at least three independent experiments. In **c** data are mean +/- s.d. from three independent experiments and each data point represents the mean of technical duplicates. Data were log-transformed and assessed for normality using the Shapiro-Wilk Test. Multiple comparisons between all groups were performed using an ordinary one-way ANOVA with Tukey’s multiple comparison test (α = 0.05). Data in **e** are from three independently infected tissue explants from 6 biological donors (n = 18). These data were log-transformed and did not meet the assumption of normality (Shapiro–Wilk test). Boxes represent the interquartile range, the line indicates the median, and whiskers show the full data range. Statistical significance between virus groups was determined using the Kruskal–Wallis test followed by Dunn’s multiple comparisons. P-values in bold indicate significance.

All the viruses tested in this study possess residue 631M in PB2 except for r-Bovine-B3.13 (Fig 5a). To further probe the role of PB2 M631L for avian virus replication in bovine cells, we introduced M631L into a selection of our 2:6 avian-origin recombinant viruses, including r-EA-2020-C, r-EA-2021-AB, r-EA-2022-BB, r-Avian-B1.1 and r-Avian-B2.1, and compared them to a r-Bovine-B3.13 (and a L631M reversion mutant) in bovine cells and in an *ex vivo* udder tissue system (Fig 5d-e). L631M reduced r-Bovine-B3.13 replication in BSF cells, as expected (Fig 5d). PB2-M631L enhanced replication of EA-2022-BB and r-Avian-B1.1 in BSF cells but did not significantly improve replication of r-EA-2020-C, r-EA-2021-AB, or r-Avian-B2.1, suggesting that the genomic context of this mammalian-adaptation is important. In *ex vivo* udder tissue, the replication pattern of M631L-viruses was similar to that observed in BSF, although no significant differences were observed for any of the viruses tested (r-Goose-B3.13, r-EA-2020-C, r-EA-2022-BB, or r-Avian-B1.1; Fig 5e). This is likely due to variability between explants in the ex vivo udder system, which makes it less suitable for detecting relatively small differences in replication efficiency (Fig S3).

We also assessed additional adaptive PB2 mutations that have arisen *in vivo* in experimentally infected cows^81^. In a recent study, the PB2 E627K mutation was selected *in vivo* in experimentally inoculated dairy cattle infected with high titres of an H5N1 2.3.4.4b of the euDG genotype^81^. We first compared the activity of the euDG polymerase with and without the E627K substitution. In human cells, polymerase activity was boosted significantly by this substitution, to levels far higher than the native euDG and Bovine-B3.13 polymerases (Fig 5f). Next, we compared the 2:6 euDG virus against the PB2-627K bearing mutant and observed a clear boost in bovine cells (Fig 5g). However, euDG-PB2-627K did not replicate to the same levels as the r-Bovine-B3.13 virus in BSF cells, further supporting the notion that a single PB2 mutation, that boosts activity in human cells, is not necessarily sufficient to fully recapitulate the replication phenotype of bovine B3.13.

Next, we assessed whether additional mutations acquired by the B3.13 virus during its evolution in dairy cattle throughout the outbreak further enhanced replication in bovine cells. We introduced a series of mutations in the internal genomic segments of our B3.13 Texas bovine virus (sampled 20^th^ March 2024) that accounted for changes that had been acquired later in the outbreaks in Colorado and California (after 3 and 5 months, respectively, of continuous circulation in cows) (Fig S6). We observed no differences in the replication of WT or mutant r-Bovine-B3.13 viruses possessing PB2-E249G and NS1-R21Q (Colorado isolate) or PB2-K670R, PA-V432I and NS1-R67G (California isolate) in bovine cells.

### Sensitivity of 2.3.4.4b H5N1 to MX1 and BTN3A3

The host type-I/III interferon (IFN) response is an important barrier to virus cross-species transmission^57,62^. Upon infection, cells sense pathogen associated molecular patterns and initiate a signalling cascade resulting in the secretion of IFNs. These IFNs, in turn, induce an antiviral state in both infected and uninfected bystander cells, by stimulating the expression of hundreds of ISGs, some of which have direct antiviral activity and can inhibit virus replication, thereby limiting within-host spread in infected tissue, pathogenicity, and transmission^82–84^. We next assessed key ISGs known to preferentially restrict avian IAVs. MX1 (also known as MXA) (dynamin-like GTPase myxovirus-resistance Gene 1), targets viral ribonucleoproteins (vRNPs)^57,59–62^ and restricts avian-origin viruses that do not possess escape mutations. A recent report showed that human MxA and bovine MX1 are both effective at restricting a bovine B3.13 virus^85^. We expanded on this work by testing a broader diversity of IAVs in both human and bovine cells backgrounds. We confirmed that MX1 is expressed in bovine udders following infection of precision cut bovine udder slices with r-Tx/37-B3.13 and examining MX1 and NP staining by immunohistochemistry and immunofluorescence (Fig 6a-b).

**Fig 6.**
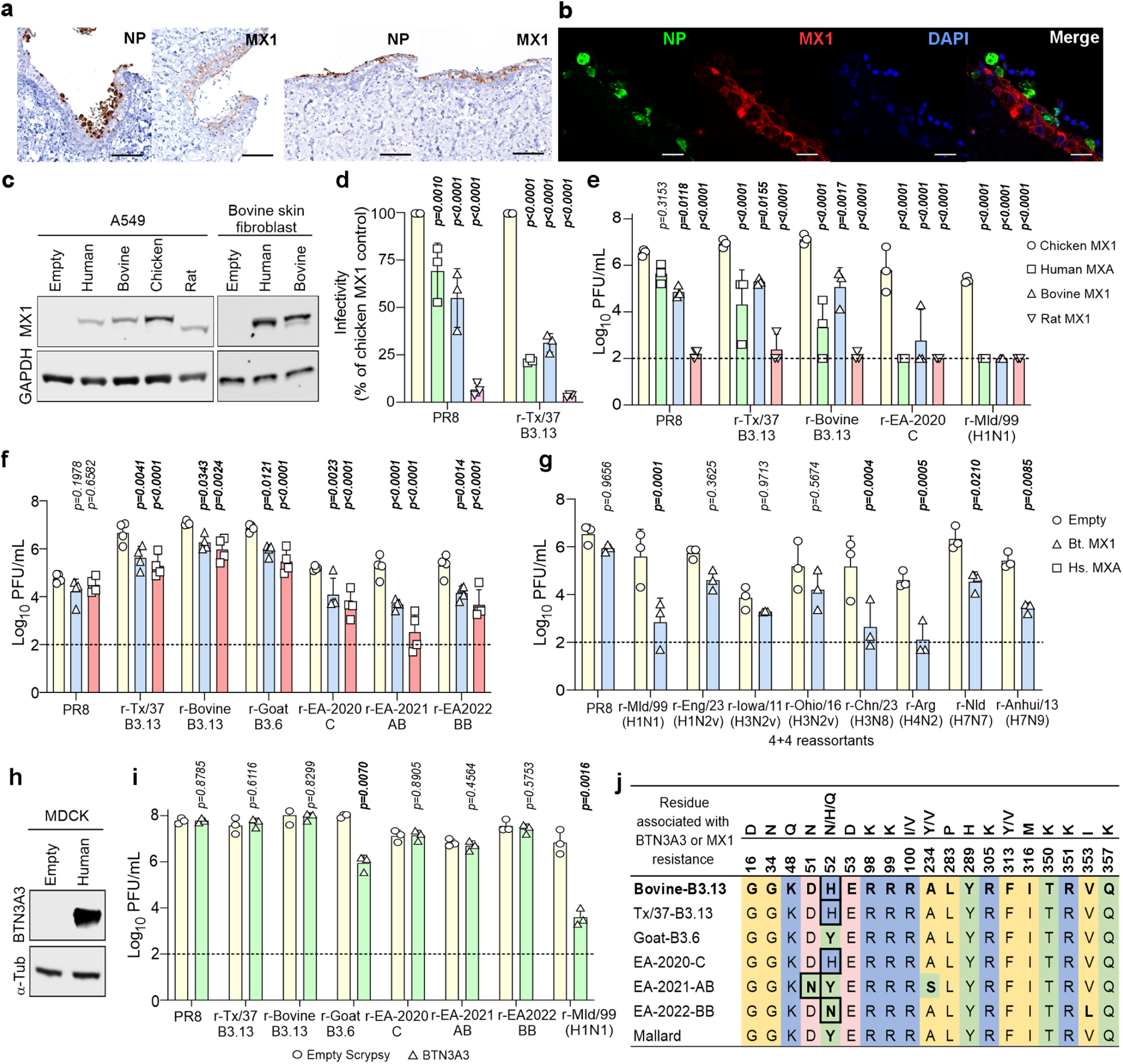
Effect of MX1 and BTN3A3 on 2.3.4.4b reassortant viruses. **a**, Udder slices infected with r-Tx/37-B3.13. Signal (brown) for NP is visible in upper the epithelial layer lining the duct, while MX1 signal is localized in the cytoplasm of ductal epithelial cells. Scale bars, 100 μm. **b**, Confocal images of udder slices stained for NP and MX1 show miminal colocalization in the duct epithelium. Scale bars, 20 μm. **c**, Western blot of lysates from A549 and BSF cells stably expressing human (MXA), bovine, chicken and rat MX1. **d**, Single-cycle titration of ZsGreen-expressing reporter viruses on modified A549 cells at 7 hpi. Titres are normalised to Chicken MX1 control cells. **e**, Modified A549 cells overexpressing MX1 were infected at MOI 0.001; infectious titres were determined at 48 hpi by plaque assay on MDCK cells. **f**, Modified BSF overexpressing MX1 were infected at an MOI of 0.001; infectious titres in the supernatant at 24 hpi were determined by plaque assay on MDCK cells. **g**, As described in **(f)** except recombinant 4:4 viruses (Table S2) were used and peak titre of each virus (24h or 48h) are shown. **h**, Western blot of lysates from MDCK cells overexpressing human BTN3A3. **i**, Virus titres on modified MDCK cells overexpressing human BTN3A3 or transduced with an empty vector control. **j**, Alignment of NP residues at positions associated with MX1 or BTN3A3 resistance. Residues are aligned to r-Bovine-B3.13 (bold). Changes are highlighted in bold; residues associated with resistance are circled in black. The experiments described are from three biological repeats, except for **(f)**, which are from four repeats. Data in **(e-h)** were log-transformed and statistical significances between groups in each experiment were measured by two-way ANOVA with Tukey’s **(d,e,f)** or Šídák **(g, i)** multiple comparisons tests. Only comparisons relative to untreated or chicken MX1 controls are shown. P-values in bold indicate statistical significance.

MX1 acts on the viral NP^59^. As our 2:6 reassortants carry the NP of bovine influenza virus (and viruses of interest) they are a suitable model to evaluate the susceptibility of different IAVs to MX1 restriction. To this end, we constitutively expressed human, bovine or rat MX1 in bovine fibroblasts and human A549 cells (Fig 6c). In human A549 cells, both bovine and human MX1 displayed antiviral activity against an r-Tx/37-B3.13 reporter virus in a single-cycle infection relative to a chicken MX1 control gene (lacking antiviral activity^86–88^) by approximately 3-fold, while PR8 was affected by less than two-fold relative to the chicken MX1 control cells (Fig 6d and Fig S7). Neither bovine or human MX1 proteins displayed the strength of antiviral activity exhibited by the rat MX1 control, a known potent antiviral gene against IAV^89^. In a low MOI, multicycle assay, both human and bovine MX1 exhibited antiviral activity against the B3.13 reassortants. However, the restriction of these viruses was less pronounced when compared to the avian viruses r-EA-2020-C or r-Mld/99(H1N1) (Fig 6e). Furthermore, both human and bovine MX1 proteins retained their antiviral properties against a wide array of both 2:6 and 4:4 reassortants when overexpressed in bovine skin fibroblasts (Fig 6f-g) suggesting that these proteins can function in both the human and bovine cellular contexts. All reassortants based on 2.3.4.4b H5N1 were restricted by both bovine and human MX1. As expected, all reassortants derived from avian viruses were inhibited by bovine MX1 (Fig 6f). In contrast, PR8 and reassortants originating from zoonotic swine viruses, r-Eng/23(H1N2v), r-Iowa/11(H3N2v) and r-Ohio/16(H3N2v), were not significantly restricted by bovine MX1 (Fig 6g). Hence, these results reinforce the notion that bovine MX1 is an effective IAV restriction factor, as has been reported recently^85^, but also highlight that different IAVs vary in their susceptibility to this restriction.

Various residues in NP that dictate Mx1/MxA sensitivity/resistance have been reported previously^60,90–92,93^. Of the 2.3.4.4b recombinant viruses tested, only r-EA-2022-BB had any of the above resistance mutations (Y52N) (Fig 6j), however this was not sufficient to overcome restriction by either human or bovine Mx1 proteins. Furthermore, the Y52H mutation, which has been described to help evade equine Mx1^93^, did not permit r-Bovine-B3.13 to evade human or bovine Mx1 proteins. This may suggest that either the viral context is important, or that bovine and human Mx1 proteins target different combinations of NP residues. From the more diverse panel of IAVs tested, the swine-origin human spillover viruses (r-Eng/23, r-Iowa/11 and r-Ohio/16) had all acquired R100I or R100V, and r-Eng/23 had also acquired F313V, and this might explain their relative resistance to bovine Mx1 (Fig 6j).

Another ISG that specifically restricts most avian viruses, and also acts on the viral NP, is BTN3A3^57^. While MX1^58–62^ is conserved in most vertebrates, including cattle, BTN3A3 originated specifically during the evolution of old world monkeys^57^, and therefore cannot play a role in H5N1 adaptation to cow cells. However, this restriction factor is a human genetic barrier to avian IAV zoonotic transmission. Hence, we assessed the replication of a selection of our reassortant viruses, including PR8 and r-Mld/99(H1N1) as controls, in MDCK cells overexpressing BTN3A3, and in the parental cells (Fig 6h-i). As expected, PR8 replicated equally well in parental and BTN3A3 expressing cells, while r-Mld/99(H1N1) replication was substantially inhibited in the latter. Of the reassortant viruses tested, only r-Goat-B3.6 was restricted by BTN3A3, while all the other 2:6-based 2.3.4.4b or B3.13 viruses evaded this restriction factor (Fig 6i).

For BTN3A3, the residues involved in sensitivity/resistance described to date include sites 52 & 313 (with a combination of 52Y & 313F/L being a sensitive genotype and one of 52H/N/Q and 313V/Y being sufficient to provide resistance). The resistant phenotype, therefore, was not unexpected in the cases of r-Bovine-B3.13, r-Tx/37-B3.13, r-EA-2020-C, and r-EA-2022-BB, which each have either NP-52H or NP-52N^57^ (Fig 6i & j). Likewise, r-Goat-B3.6 and r-Mld/99 both contained the sensitive genotype of NP-52Y and NP-313F^57^. Interestingly, however, r-EA-2021-AB was resistant to BTN3A3 activity, despite having NP-52Y & NP-313F, suggesting that other residue(s) are important. Hence, many 2.3.4.4b viruses already possess NP mutations that evade BTN3A3 restriction. We note that r-Goose-B3.13 was not tested in this series of experiments because its NP is identical to r-Bovine-B3.13.

### Increased modulation of the type-I IFN response by bovine-B3.13

Above we showed that bovine MX1 possess antiviral properties and both avian and bovine-adapted viruses are restricted by this ISG. As highlighted above, expression of MX1, and other antiviral ISGs, is under the control of the host the type-I/III IFN response, which is therefore in its complexity an important barrier to virus cross-species transmission^57,62^. We next compared the susceptibility of r-Bovine-B3.13, r-Tx/37-B3.13, and r-EA-2020-C to the bovine type-I IFN response. First, we confirmed that stimulation of bovine fibroblasts with universal type-I IFN resulted in the activation of markers of a type-I IFN response. As expected, western blot analysis of IFN-treated bovine fibroblasts showed a dose-dependent upregulation of phosphorylated Stat1 (pSTAT1), and of the ISGs MX1 and RSAD2 (Fig. 7a). Next, we compared the susceptibility of r-Bovine-B3.13, r-Tx/37-B3.13, and r-EA-2020-C to IFN by assessing their replication in pre-stimulated or mock-treated bovine fibroblasts. All viruses showed sensitivity to IFN in a dose-dependent manner (Fig 7b). However, r-EA-2020-C displayed a greater susceptibility to the bovine IFN response than the B3.13 viruses, or PR8 used as control. For example, at 24 hpi, pre-stimulation of bovine fibroblasts with 40U/ml of IFN was sufficient to completely inhibit replication of r-EA-2020-C below the detection limit, while both r-Bovine-B3.13 and r-Tx/37-B3.13 reached titres around 10^4^ PFU/ml in the same conditions. At 48 hpi, 1.6 U/ml of IFN had no effect on r-Bovine-B3.13 titres, but reduced r-EA-2020-C titres 150-fold on average (Fig. 7b, right). In addition, the avian B3.13-derived r-Goose-B3.13 was as similarly sensitive to IFN as r-EA-2020-C (Fig. 7c-d).

**Fig 7.**
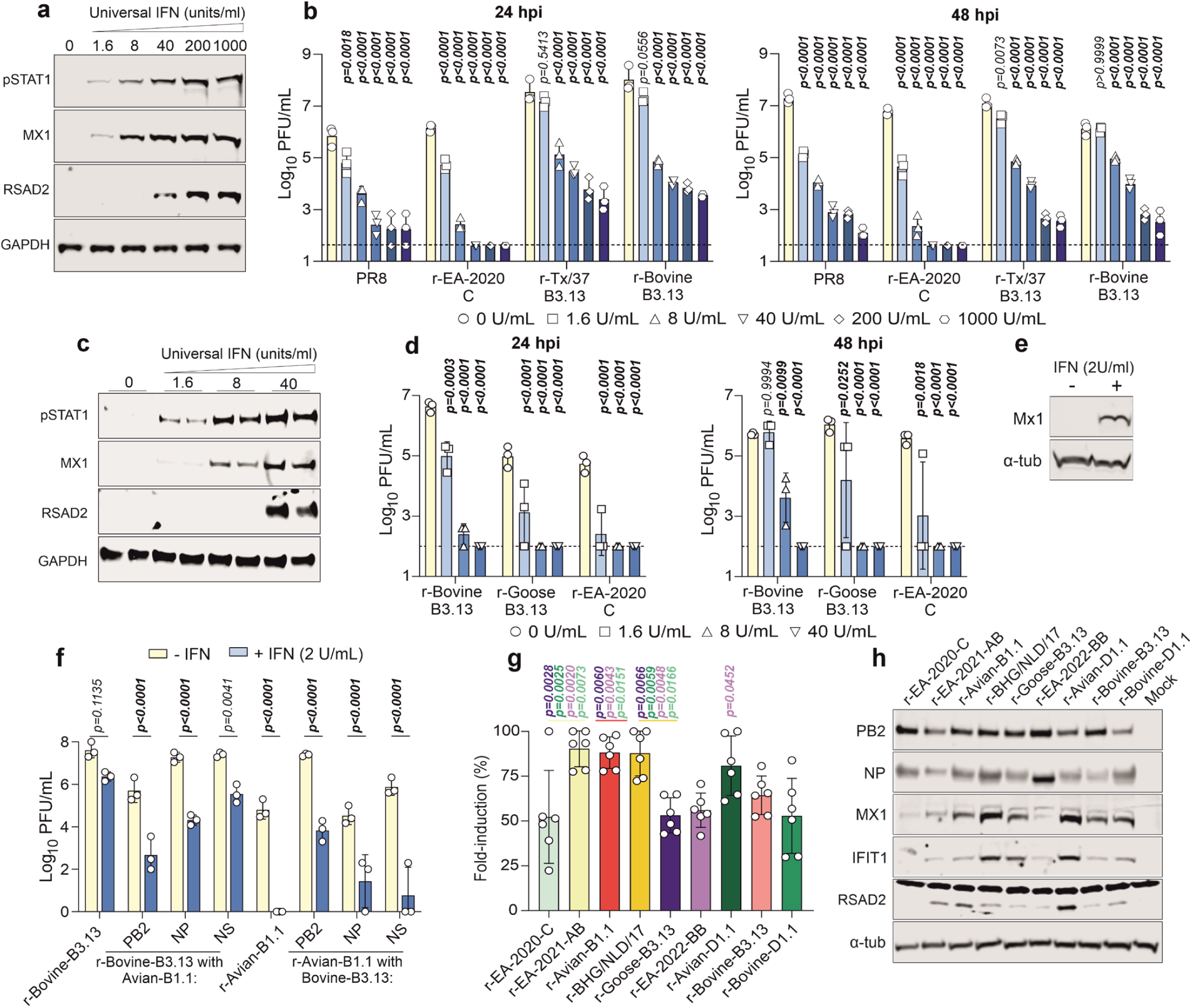
Sensitivity of 2.3.4.4b reassortant viruses to type I IFN in bovine cells. Immortalised BSF were pre-treated for 24 h with increasing concentrations of type I IFN prior to low MOI infection (0.001 PFU/cell). **a**, Western blot of interferon-stimulated genes (pSTAT1, Mx1, RSAD2) at 24 h post IFN-treatment. **b**, Virus titres in supernatants were determined by plaque assay on MDCK cells at indicated time points post-infection. **c,d**, As in **a** and **b** but including r-Goose-B3.13. **e-f**, Replication of r-Bovine-B3.13, r-Avian-B1.1 and reciprocal segment swaps in the absence or presence of universal IFN. BSF were pre- or mock-treated with 2 U/mL IFN prior to infection at MOI of 0.001. Western blot for MX1 at 24 hours post treatment **(e)**; viral titers at 24 hpi were determined by plaque assay on MDCK cells **(f). g**, Immortalized udder skin fibroblasts expressing an ISRE-firefly luciferase reporter were infected with 2.3.4.4b 2:6 viruses at an MOI of 5 PFU/cell. Luminescence was measured at 24 hpi and normalised to mock controls. **h**, Western blot of viral proteins and ISGs in udder fibroblasts infected at an MOI of 5 for 24 hours. Data represent means from three biological replicates (average of two technical replicates per data point); error bars show +/- s.d. Data in **g** are from six biological replicates. For **b-f** data were log-transformed; statistical significances in **b-f** between IFN treated and untreated groups was assessed by two-way ANOVA with Dunnett’s (**b-e**) or Šídák **(f)** multiple comparisons tests (α = 0.05). For **g**, normality was confirmed by Shapiro-Wilk test; comparisons were determined by one-way ANOVA with Tukey’s multiple comparisons. For **b-f**, P values in bold indicate statistical significance. For **g**, only significant values are shown.

To attempt to dissect this phenotype, we infected BSF pre-treated with 2 U/mL of IFN with some individual segment swaps between r-Bovine-B3.13 and r-Avian-B1.1, the latter representing one of the 2.3.4.4b genotypes in which PB2 M631L enhanced viral replication in bovine cells (Fig 7e-f). We generated reassortants with segments 1, 5 and 8 (encoding PB2, NP and NS1/2) because B1.1 with bovine B3.13 PB2 provides an opportunity to assess an otherwise avian virus that replicates similarly to parental r-Bovine-B3.13 in the absence of IFN (Fig 4g). In addition, NP is a target of Mx1 (and BTN3A3)^57–60^, and NS1 is a known viral modulator of the IFN response^94,95^. This experiment revealed that the parental r-Bovine-B3.13. was the least sensitive to 2 U/mL IFN treatment, as each of the avian B1.1 PB2, NP, and NS reassortants showed reduced virus replication in the face of an IFN response (Fig 7f). Importantly, in the absence of IFN, r-Bovine-B3.13 with B1.1 NP or NS segment swaps and r-Avian-B1.1 with the Bovine-B3.13 PB2 replicated similarly to the parental r-Bovine-B3.13 virus. These data suggests that the bovine-B3.13 internal genes are better equipped to deal with an IFN antiviral state in bovine cells than the equivalent B1.1 counterpart genes, and highlighting a role for the PB2, NP and NS1.

We next measured the ability of diverse recombinant 2:6 viruses to modulate the interferon response using primary bovine udder fibroblasts expressing a luciferase gene under the control of an interferon response element (ISRE) in bovine udder fibroblasts. We infected ISRE-luciferase reporter cells at MOI 5 PFU/cell and measured the induction of ISRE-driven luciferase. All viruses tested induced ISRE-dependent luciferase expression relative to mock-infected cells, although the fold-induction varied across the viruses tested. Viruses that were better at supressing ISRE-luciferase included r-EA-2020-C, r-Goose-B3.13, r-EA-2022-BB, r-Bovine-B3.13 and r-Bovine-D1.1 (Fig 7g-h). In comparison, r-EA-2021-AB, r-AvianB1.1, r-BHG/NLD/17 and r-Avian-D1.1 were less efficient at supressing ISRE-luciferase expression. These data highlight the varied ability of genetically diverse H5N1 to supress ISG expression in infected bovine cells.

To complement these experiments, we also tested a subset of viruses for their ability to induce host protein shutoff by measuring incorporation of puromycin into nascent proteins as a marker for efficient cellular protein synthesis. To this end, we infected BSF with a selection of reassortants at MOI 5 for 16h or 24h, then chased with puromycin for one hour before harvest. Our results (Fig. S8) shows that the ruminant-derived recombinant viruses (r-Bovine-B3.13 and r-Tx37-B3.13) and the avian r-EA-2022-BB induced a more severe cellular shutoff than r-EA-2020-C and r-Goose-B3.13, in line with their replication ability in these cells.

Collectively these results suggest that the ability to modulate the host IFN response also varies between 2.3.4.4b bovine-adapted and avian genotypes, with important differences, particularly among the latter.

## DISCUSSION

This study shows the multiple determinants of IAV host range and suggests that the replication fitness of H5N1 viruses in bovine cells varied during their evolutionary history (Fig. 8). H5N1 has been considered a threat to global health since the emergence of the Gs/Gd lineage in poultry in 1996, with the associated spillovers and fatal cases in humans. While numerous spillover events of avian IAV (including H5N1) from birds to humans and other mammals have been reported over the years prior to the most recent dairy cattle outbreaks, only a few avian viruses have established themselves as endemic mammalian lineages (Eurasian H1N1 in pigs^96^, H7N7 and H3N8 in horses^97–99^, and H3N2 in dogs^100^).

**Fig 8.**
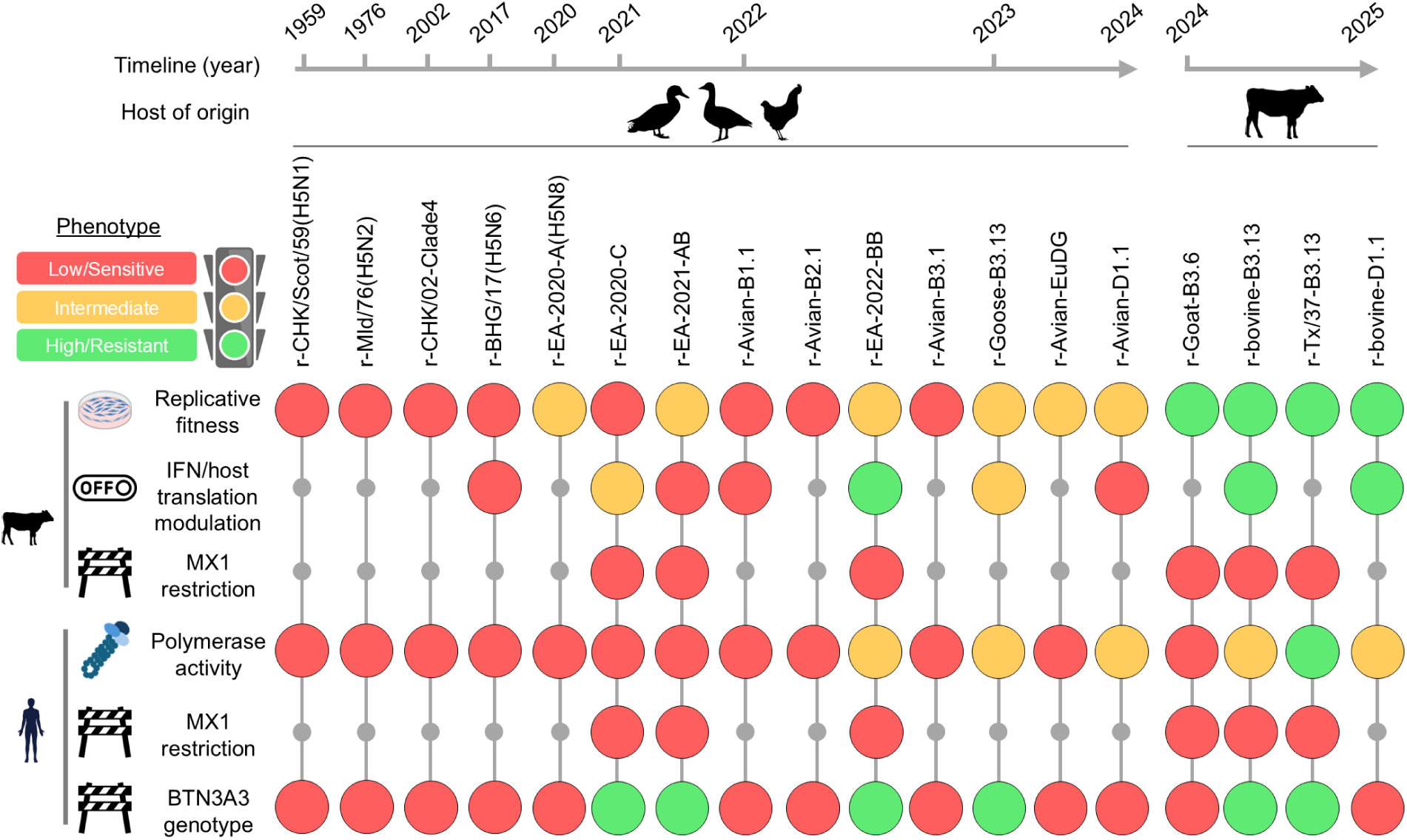
Phenotypes of the reassortant viruses observed in this study. Schematic diagram summarising the phenotypes of the reassortant viruses described in this study using either bovine or human cells and restriction factors as indicated. Note that summary of “IFN/host translational modulation” derives from results shown in Fig 7 and S8.

The global emergence of the 2.3.4.4b clade has reignited justifiable concerns about the possibility of an H5N1 pandemic. Both the geographical distribution and the number of mammalian species infected by clade 2.3.4.4b viruses have dramatically increased compared to past H5N1 infection waves^7^. In addition, the current epizootic in dairy cattle provides the clearest example of a direct transfer of an entire avian IAV into a new mammalian host^101^. Importantly, mammal-to-mammal transmission of H5N1 2.3.4.4b has occurred also in other phylogenetically distant species such as marine mammals in South America^14,16,17,102^, and farmed fur animals in Europe^103,104^.

While it is likely that the diversity of animal species affected by H5N1 reflects in part the increased geographical expansion of this virus^101^, we reasoned that the 2.3.4.4b clade may also possess intrinsic biological properties favouring spillover in mammals. The viral HA (which defines the 2.3.4.4b clade), and its ability to bind host cell receptors, is a well-recognised determinant of IAV host tropism and several studies on B3.13 have already focused on this aspect^38–47^. However, within the 2.3.4.4b clade, there are more than 80 genotypes determined by their internal gene cassette.

Our study indicates that the constellation of internal genomic segments of the B3.13 and D1.1 viruses established in cows (including also the zoonotic human B3.13 case in Texas and the B3.6 genotype that caused an outbreak in goats) provides a clear replication advantage in bovine cells over all the other IAV we tested, including related avian 2.3.4.4b viruses and other IAVs originating from humans, pigs, horses and dogs. In addition, this phenotype is not confined only to cells from the udder, considering we tested additional bovine cell types. Hence, while HA-receptor preference and high virus titres in the milk and farming practices clearly favoured the spread of H5N1 in udders of dairy cows, the internal gene cassette of viruses such as B3.13 and D1.1 can potentially replicate in more than one type of bovine cells.

Importantly, we also show that the replicative fitness of avian H5N1 viruses in bovine cells varied greatly. Reassortants with internal gene cassettes belonging to genotypes that spillover in dairy cow, such as avian B3.13, avian D1.1 and those spilling over in fur farm mammals, such EA-2022-BB ^73,104^, displayed the highest replication efficiency, along with others including EA-2020-A, and euDG. Interestingly, the latter was shown to be able to replicate in the udder of dairy cows experimentally infected via the intramammary route with high virus load^81^. Others, such as recombinants based on EA-2020-C, the North American B3.1, B2.1, B1.1 and an early 2.3.4.4b virus isolated in 2017 in the Netherlands reached significantly lower titres in bovine cells.

Overall, our data suggest that H5N1 viruses possessing genetic traits that confer bovine fitness are circulating in Europe and North America, and viral emergence might be impaired by other barriers such as agricultural practices. The recent detection of H5N1 in a sheep in the UK^105^ provides a tangible example of the global risk that these viruses pose to ruminants regardless of whether the H5N1 outbreaks in North America are eventually controlled. The replication fitness in bovine cells may be a phenotype acquired with the evolution of the 2.3.4.4b clade. Indeed, our reassortants possessing the internal gene cassette from H5N1 viruses isolated before the expansion of the 2.3.4.4b clade, including viruses isolated between 1959 to 2002, replicated particularly poorly in bovine cells.

Using single and multiple segment swaps between recombinant viruses, our results suggest that the European avian 2.3.4.4b genotype (EA-2020-C) thought to be the one that originally moved into North America from Europe, requires various, simultaneous adaptive mutations in multiple segments of the polymerase complex and possibly NS, to replicate efficiently in bovine cells. Indeed, PB2, PB1, NP and NS were all required to switch the phenotype of the ancestral European r-EA-2020-C in bovine cells. Conversely, only the PB2 of a recombinant virus representing the closest sequenced American avian B3.13 predating the epizootic in cattle (r-Goose-B3.13), and for a distinct B1.1 genotype, was sufficient to switch the phenotype of r-Bovine-B3.13. Importantly, three non-synonymous mutations in PB2 differentiate r-Goose-B3.13 from r-Bovine-B3.13 (see below). Among these, the M631L mutation has been already shown to increase viral replication in mammalian cells^7,71,72^. Our data however suggest that the phenotype provided by this mutation is context dependent, as it does not boost replication of all the 2.3.4.4b viruses that we tested, and that in particular PB2 441N may play an additive role in boosting replication. Interestingly, this mutation arose independently in a ferret transmission study within a neuraminidase-inhibitor resistant variant of the H7N9 virus that caused extensive human spillovers^106^. Further, while this mutation was not widespread in avian B3.13 viruses closely related to the bovine-B3.13 virus^22^, it has been maintained in sub-clades of B3.13 bovine viruses (Fig S6).

Of note, H5N1 A/Texas/37 (a B3.13 virus isolated from a human spillover case) acquired the well-described PB2-E627K mutation, a relatively frequent avian-to-mammalian mutation known to enhance interactions with human ANP32 proteins^50,107^. The E627K mutation is instead absent in the PB2 of both B3.13 and D1.1 bovine H5N1 (while the latter has the well-known D701N). Interestingly, using a panel of 33 distinct IAV polymerases in replicon assays, we showed that those carrying PB2 627K, including the human B3.13 H5N1 Tx/37, displayed the highest activity in human cells, significantly higher than the bovine- and goat-derived H5N1 polymerases. These are all polymerases derived from viruses that are either mammalian-adapted or derived from spillover events in humans. Our data in bovine cells using 4:4 reassortants with the polymerase complex of the viruses of interest, supported some of the differences noted above. Thus, overall, our data show clearly that adaptation of B3.13 and D1.1 (and B3.6), is bovine-specific and distinct from a broader polymerase adaptation to mammalian cells that could be detected in human cells. Consistent with this notion, recombinant viruses possessing the internal genes of other mammalian-adapted viruses did not replicate as efficiently in bovine cells as those with bovine B3.13, D1.1 and goat B3.6 internal genes. Interestingly, in human cells there were little or no differences between the polymerase activity of the bovine-adapted H5N1 viruses and those of avian viruses from genotypes that have spilled over in bovine or fur animals (avian B3.13, D1.1 and EA-2022-BB). However, activities of this group of polymerases were higher than most other remaining avian polymerases. In mice experiments, r-Bovine-B3.13 caused more severe infection than avian r-EA-2020-C virus.

We found the bovine and human B3.13 reassortant viruses were also less sensitive to the bovine type-I IFN response compared to avian EA-2020-C, an avian B3.13 ancestor, and an avian B1.1 virus. H5N1 B3.13 were restricted by bovine (and human) MX1, and human BTN3A3, two known restriction factors blocking avian IAV replication^58–62^. When we supplied an avian B1.1 virus with a bovine B3.13 PB2, so that replication in the absence of IFN was comparable to the bovine B3.13 equivalent, we observed a greater sensitivity to IFN in bovine cells. Similarly, when the bovine NP and NS segments were replaced with the avian B1.1 equivalent, this also increased IFN sensitivity, indicating adaptive roles for NP and NS in counteracting the IFN response. We found the bovine and human B3.13 reassortant viruses were also less sensitive to the bovine type-I IFN response compared to avian EA-2020-C, an avian B3.13 ancestor, and an avian B1.1 virus. H5N1 B3.13 were restricted by bovine (and human) MX1, but not human BTN3A3, two known restriction factors blocking avian IAV replication^58–62^. When we supplied an avian B1.1 virus with a bovine B3.13 PB2, so that replication in the absence of IFN was comparable to the bovine B3.13 equivalent, we observed a greater sensitivity to IFN in bovine cells. Similarly, when the bovine NP and NS segments were replaced with the avian B1.1 equivalent, this also increased IFN sensitivity, indicating adaptive roles for NP and NS in counteracting the IFN response.

Overall, bovine-adapted viruses displayed a better ability to modulate IFN. By puromycin assays, we show that reassortants carrying the internal genes of bovine-adapted B3.13 viruses induce stronger cellular shutdown, than avian B3.13 or EA-2020-C. Interestingly, EA-2022-BB induced a shutdown as strong as the bovine-adapted viruses. pStat1-inhibition, a marker of the IFN-response activation, was also directly correlated to the observed shutdown, and we obtained similar results in cells with a reporter gene under the control of the ISRE.

We acknowledge limitations of our study. We specifically focused on the internal genomic segments of B3.13 and D1.1 and therefore have used recombinant viruses harbouring the HA and NA of the laboratory-adapted PR8. Consequently, cells that may not be permissive to a recombinant virus harbouring the glycoproteins of PR8 may be susceptible to infection by an authentic 2.3.4.4b H5N1 virus and *vice versa*. For this reason, we used immortalised bovine skin fibroblasts and primary nasal fibroblasts as cell model systems. These cells allowed us to explore species-specific virus-host interactions within a bovine intracellular context as they express the required cellular co-factors, they are IFN-competent and express known IAV-restriction factors (e.g. MX1). We selected key 2:6 viruses representing substantial IAV diversity and confirmed the phenotypes observed in these fibroblastic cells were recapitulated in *ex vivo* udder tissue from multiple animals. In addition, while reassortants are exceedingly useful tools to map molecular determinants of specific phenotypes, they may possess unforeseen defects due to incompatibilities between genome segments that have not co-evolved together. However, to mitigate for this risk, we always assessed the capacity of reassortants to replicate in chicken fibroblast cells (DF-1).

We showed that the NP of bovine-adapted 2.3.4.4b viruses can escape human BTN3A3 restriction^57^, one of the human genetic barriers to avian IAVs. Thus, the spread of 2.3.4.4b viruses in several mammalian species, including its recent adaptation to cows, increases the pool of IAVs carrying genetic traits that counteract an important human barrier against zoonotic influenza, providing abundant ecological opportunities for spillover infections that might in turn facilitate adaption to humans^108^. For example, bovine MX1 may provide a selective pressure on bovine H5N1 to select mutants that could increasingly escape both the bovine and human orthologues of this key restriction factor. These considerations, in addition to recent reports showing that a single amino acid residue in the viral HA is sufficient to switch its tropism from avian to mammalian receptors^44,45^, suggest that the zoonotic potential of H5N1 2.3.4.4b is relatively high.

## METHODS

### Data Availability Statement

Raw data underpinning the figures associated with this manuscript are available in the open access Enlighten repository (https://doi.org/XXX).

### Ethical statement

All experiments were approved by the local genetic manipulation safety committee at the University of Glasgow (GM223), and the Health and Safety Executive of the United Kingdom. Reassortants derived by reverse genetics were made with the external glycoproteins of the attenuated vaccine strain A/Puerto Rico/8/1934 (H1N1)^109^ which is mouse adapted and attenuated in humans^110,111^. The PR8 strain used in this study also possesses a strong receptor preference for avian-type α2-3 linked sialic acid^112^. Work with reassortants used in this study was physically segregated from work with mammalian viruses with glycoproteins different from PR8. All animal work was in accordance with the animal ethics and welfare committee at the University of Glasgow and the United Kingdom Home Office regulations (ASPA, 1986, PPL PP4085778). Work using animal tissues was approved by the Ethics Committee of the School of Veterinary Medicine of the University of Glasgow (ethics approval EA26/25).

### Phylogenetic analysis

Influenza A virus sequences were retrieved from the NCBI Entrez databases using the Influenza A virus taxonomic ID with an in-house Python tool to access the E-utilities API. This dataset was further enriched with sequences from GISAID matching the search term “Clade=2.3.4.4b” and consensus genomes available in Andersen Lab’s public GitHub repository (https://github.com/andersen-lab/avian-influenza). All databases were last accessed on July 31^st^, 2025. Metadata associated with the sequences available on GISAID used in this study can be accessed via EPI_SET_250912fu (DOI: https://doi.org/10.55876/gis8.250912fu). Fasta sequences were sorted by genomic segment and redundancies between databases were removed based on the name of the isolate. The MMseqs2 software tool v15.6f452 was used to cluster the sequences based on a 0.95 identity threshold and select a representative sequence per cluster (--cluster-mode 2 --cov-mode 1 -- min-seq-id 0.95)^113^, resulting in the following number per genomic segment: PB2: 458, PB1: 406, PA: 405, NP: 221, MP: 112, NS: 222. Representative sequences per segment along with the sequences of the viruses of interest (detailed in Table S1) were aligned using MAFFT v7.453 using the default parameters^114^. Maximum likelihood trees were obtained with IQ-tree v2.1.2 under the best-fit model and plotted in RStudio using ggtree v3.12.0^115,116^.

### Generation of reassortant viruses harbouring PR8 HA and NA glycoproteins

pHW2000 reverse genetics plasmids used to rescue recombinant viruses based on A/California/04-061-MA/2009(H1N1) virus were a kind gift of Prof. Daniel Perez, the A/mallard/Netherlands/10-Cam/99(H1N1) virus from Prof. Laurence Tiley, A/Puerto Rico/8/34 (H1N1)^109^, and the A/canine/Illinois/11613/2015 (H3N2) from Prof. Luis Martinez-Sobrido. Each internal gene segment from A/equine/South Africa/4/2003(H3N8) was subcloned into pHW2000 following RT-PCR from RNA extracted from virus stocks. pHW2000 reverse genetics plasmids to rescue the remaining viruses described in Tables S1 and S2 were synthesised and cloned by GeneArt Thermo Fisher. pHW2000 plasmids to generate r-Goose-B3.13 were generated by site directed mutagenesis of r-Bovine-B3.13 by introducing in the latter non-synonymous mutations in PB2, PB1, PA, and NS1/NS2 to obtain genomic segment expressing viral proteins identical to A/goose/Colorado/2024, GISAID accession EPI_ISL_19228459.

2:6 viruses with the HA and NA genes from PR8 (Table S1) or 4:4 with the HA, NA, M and NS from PR8 (Table S2) were rescued by reverse genetics. HEK-293T (or cells modified with a lentiviral vector to overexpress *Gallus gallus* ANP32A; 293T-Gg.ANP32A. Annotated in Tables S1 and S2) cells were transfected with pHW2000 reverse genetics plasmids (250 ng of each plasmid in 6-well plate format) and the media changed to ‘virus growth medium’ (serum-free DMEM supplemented with 1 ug/mL TPCK-trypsin (Sigma) and 0.14% (w/v) BSA fraction V (Gibco) at 16 h post-transfection. Supernatant was collected at 48h post-transfection and used to inoculate cultures of MDCK (or MDCK cells modified with a lentiviral vector to overexpress Gg.ANP32A; MDCK-Gg.ANP32A. Annotated in Tables S1 and S2) cells maintained in virus growth medium until approximately 70-80% of cells displayed cytopathic effect. At this point the supernatant was clarified by centrifugation prior to storage at –80 °C in small volume aliquots. All infections were carried out with previously unthawed virus aliquots to avoid freeze-thaw changes to infectious titre. All viruses used have been sequenced to confirm identity and identify spurious mutations.

### Cells

Cells were maintained in DMEM (Gibco 31966021) supplemented with 9% (v/v) HI-FCS (Gibco), 100 U/mL Penicillin and 100 µg/mL Streptomycin (Gibco) (hereafter referred to as ‘complete DMEM’) at 37°C with 5% CO_2_ unless otherwise stated. HEK-293T, MDCK, A549 cells (and their Gg.ANP32A-overexpressing derivatives) have been described previously^57^. Primary-derived bovine skin fibroblasts have been described previously^117^. Nasal respiratory epithelium from the turbinates of the cow were collected from the abattoir in complete DMEM media containing 10% FCS, 100 U/mL Penicillin and 100 µg/mL Streptomycin (Gibco) and 2.5 µg/mL Amphotericin B as an antibacterial and antifungal agents respectively. Tissues were washed three times with PBS and incubated in 10 ml of complete DMEM media containing collagenase (2 mg/ml, collagenase from *Clostridium histolyticum*, Sigma) at 37°C for 2 h. Subsequently, supernatant was strained with 70 μm strainer (Greiner) and subsequently centrifuged at 188 *x g* for 5 minutes. The cell pellet generated was washed three times in complete DMEM media to remove collagenase. Cells were finally cultured in complete DMEM and incubated at 37°C, 5% CO_2_. Media was changed every day initially to remove non adherent cells until an adherent population of fibroblasts was achieved; cells were subsequently passaged using standard procedures. All infection experiments were carried out with early passages cells (before stationary phase was achieved) as described below.

Fibroblasts were extracted from udder tissue slices generously provided by Prof. Dirk Werling, using a protocol like the one described above for the nasal fibroblasts. These cells were cultured in DMEM/F-12 (1:1) media (Gibco 11039021) supplemented with 9% (v/v) HI-FCS (Gibco), 100 U/mL Penicillin and 100 µg/mL Streptomycin (Gibco), 2.5 µg/mL Amphotericin B (Gibco) and 1X Insulin-Transferrin-Selenium-Ethanolamine (ITS -X) (Gibco) at 37°C with 5% CO_2_ and were immortalized as previously described^118^. To generate an ISRE-luciferase reporter cell line, immortalized bovine udder fibroblasts were transduced with a lentiviral vector (pGreenFire1-ISRE Lentivector) kindly provided by Dr. Adam Fletcher. This construct drives the expression of GFP and firefly luciferase under the control of the human interferon response elements (ISRE) paired with a minimal CMV promoter.

BTN3A3 overexpressing cell lines have been described previously^57^. To overexpress MX1, the Mx1 genes of *Rattus norvegicus* (GenBank accession NM_173096.3), *Bos taurus* (GenBank accession NM_173940.2), *Homo sapiens* (GenBank accession NM_002462.5) and *Gallus gallus* (GenBank accession NM_204609.2) were synthesised by BioBasic and subcloned into the SCPRSY lentiviral vector. Cells were modified by lentiviral transduction using SCRPSY vectors using standard techniques as previously described^119^.

### Virus replication assays

Typically, cells were plated in 12-well plates the day prior to infection. Cells were washed with serum-free DMEM prior to infecting with 200 µL of virus diluted in serum-free DMEM to achieve desired MOI (MOI stated in respective figure legends).

After 1 h of incubation, the inoculum was removed and cells were overlaid with 1 mL of virus growth medium. Standard plaque assays to titrate infectious virus in the supernatant of infected cells were performed in MDCK cells as described previously^57^.

### Quantification of viral RNA

Infected culture supernatants were homogenized in Trizol LS reagent (Thermo Fisher), and viral RNA was extracted using the Direct-zol-96 MagBead RNA kit (Zymo Research, R2102) automated on the KingFisher Flex System (Thermo Fisher) as per the manufacturers’ instructions. To quantify negative-sense vRNA and avoid non-specific priming of positive-sense replication intermediates, quantification was carried out as a two-step RT-qPCR. Specifically, vRNAs were reverse transcribed using SuperScript IV (Thermo Fisher) and the Uni12 primer^120^ in half-reactions containing 2.5 ul of extracted RNA. qPCR was performed using Fast SYBR Green Master Mix (Thermo Fisher), and HA fragment specific primers: IAV_HA_For 5ʹ AAACAGCAGTCTCCCTTTCCAG and IAV_HA_Rev: 5ʹ GTTCCTTAGTCCTGTAACCATCCTCA. Each reaction contained 4 ul of cDNA and was run using an ABI7500 Fast instrument, according to the manufacturer’s recommendations, and results were analysed with the 7500 Software v2.3 (Applied Biosystems, Life Technologies). A 714 bp synthetic gBlock carrying a partial sequence of PR8 HA (Twist Bioscience, custom) was used to generate a standard curve. Viral load was extrapolated from the curve and viral genomes were expressed as number of HA gene copies per ml of supernatant.

### Polymerase activity assays

Subconfluent monolayers of 293T cells (1.5 × 10^5^ cells seeded in 24-well plates the day prior to transfection) were co-transfected with 10 ng each of pHW2000 bi-directional plasmids encoding PB2, PB1, PA and transfection control plasmid (CMV-driven expression of Renilla luciferase) and 25 ng each of NP and a PolI-driven expression of cRNA-sense Firefly luciferase reporter RNA, in Gibco™ Opti-MEM™ Reduced Serum Medium, using Invitrogen™ Lipofectamine™ 2000 CD Transfection Reagent. Transfection lacking the NP plasmid served as negative control. 24 hours post-transfection, the transfection medium was removed, and cells were lysed by adding 100 µl of 1X-passive lysis buffer (Dual-Luciferase reporter assay system, Promega), and freeze-thawing. The lysates were transferred to white-walled 96-well plates (Greiner) and luminescence was measured from 20 μL of lysate by using 30 μL of either luciferase assay reagent II or STOP & Glo reagent (Dual-Luciferase reporter assay system, Promega) in a GloMax® Navigator Microplate Luminometer (Promega) to measure IAV polymerase-driven Firefly luciferase activity and cellular-driven Renilla luciferase activity, respectively. Firefly luciferase values for every sample were normalised by their respective Renilla luciferase values to account for transfection efficiency variation between samples. Each sample was tested in duplicate in at least in three separate biological experiments.

### MX1 and BTN3A3 susceptibility assays

For single-cycle titration of ZsGreen reporter viruses on A549-Mx1 cells, cells were cultured in 96-well plates 16 h prior to infection (1.5 × 10^4^ cells/well). Cells were washed with serum-free DMEM prior to infection with 100 µL of serum-free DMEM containing serially-diluted virus. At 7 h post-infection, cells were washed with PBS and dispersed with TryLE (Gibco), resuspended in complete DMEM and fixed in 2% formaldehyde. To measure ZsGreen-positive cells, flow cytometry was performed on a Millipore GUAVA EasyCyte HT Flow Cytometer. Percentage of ZsGreen-positive cells was determined using FlowJo (https://www.flowjo.com) software (Fig S3).

### Immunoblotting

Total cell lysates were prepared as previously described^121^. Lysates were heated at 75 °C for 10 min and subjected to polyacrylamide gel electrophoresis before being transferred to polyvinylidene difluoride (Merck Millipore IPFL00010) membranes at 30 V for 90 min. Membranes were blocked for 1 h with 1X Tris-buffered saline-0.2% Tween 20 (TBST) containing 5% dried skimmed milk, followed by incubation with primary antibodies diluted in 5% BSA/0.01% sodium azide in 1X TBST, overnight at 4 °C. After four 10-min 1X TBST washes, membranes were incubated with secondary antibodies (diluted in blocking buffer), for 1 h at room temperature protected from direct light. Following four more washes in 1X TBST, membranes were imaged using the LI-COR CLx-Odyssey Imaging platform. The primary antibodies used in this study are: PB2 - GeneTex (GTX125926), NP - MRC PPU Reagents and Services, Dundee (DA183, 5^th^ Bleed), NS1 – (MRC PPU Reagents and Services, Dundee (DA182, 2^nd^ Bleed), pSTAT1 (Tyr701) - Cell Signaling (9167S), IFIT1 - Origene (TA500948), RSAD2 - Proteintech (28089-1-AP), GAPDH - Cell Signaling (2118S), α-tubulin - Proteintech (66031-1-Ig), β-actin - Proteintech (66009-1-Ig), Mx1 - Proteintech (13750-1-AP) and puromycin (Millipore; MABE343). A Mx1 antibody raised in mice was kindly provided by Georg Kochs (University Medical Centre, Freiburg, Germany). Secondary antibodies used: Anti-rabbit IgG (H+L) (DyLight 800 4X PEG Conjugate) Cell Signaling (5151S), Anti-mouse IgG (H+L) (DyLight 680 Conjugate) Cell Signaling (5470S) and Anti-sheep IgG (H+L) (Alexa Flour^TM^ 680) Thermo Fisher Scientific (A21102).

### Precision-cut bovine udder slicing and *in vitro* culture

Precision-cut bovine udder slices (PCBUS) were prepared from udders of commercially slaughtered dairy cows without mastitis or any antibiotic treatment. Subsequently, a piece of approximately 10cm x 10cm x 5 cm was cut out of the middle region of the bovine udder, immediately transferred into Krebs-Henseleit solution and transported to the laboratory. Tissue was cut initially into slices of roughly 1 cm, and subsequently into tissue cores with a diameter of 8 mm. To stabilize the tissue in the tissue core holder, the tissue was embedded in 3% agarose (w/v in 0.9% NaCl). PCBUS were prepared with a Krumdieck tissue slicer with a thickness of 250-350 µm. The slicer was filled with ice-cold pre-oxygenated Krebs-Henseleit solution, adjusted to a pH of 7.4. Promptly, slices were placed in a 24-well plate in 1mL of pre-warmed and pre-oxygenated medium.

Incubating slices was done in RPMI-1640 medium supplemented with 1% penicillin-streptomycin, 1% fungizone and different concentrations of FCS (0%, 2% or 10%) at 38.5°C in presence of 5% CO_2_. To assess cell viability, PCBUS were examined every 24 hours until the end of the experiment using AlamarBlue®. To evaluate cell integrity, HE-staining was performed. After infection, slices were fixed in 4% formalin at 4°C for 24 h. Afterwards, slices were dehydrated in ethanol with increasing concentrations. Slices were subsequently cleared in xylene. Thereafter, slices were horizontally embedded in paraffin wax and sectioned at 2-3 μm. Prior to staining with haematoxylin and eosin (H&E), sections were deparaffinized and rehydrated. The cell viability of udder tissue was assessed by evaluating the cytoplasm and the shape/staining of nuclei as well as presence of necrosis and apoptosis.

### IFN-susceptibility assays

Immortalised bovine skin fibroblasts were seeded at a density of 2.5 × 10^5^ cells/well of a 24 well plate in complete DMEM media. Next day to immune prime the cells, media was removed and complete DMEM media containing indicated concentrations of Universal Type I IFN (Human IFN-Alpha Hybrid Protein from PBL Assay Science, Catalog Number: 11200) were added to the cells for 24 hrs. The next day, media was removed and cells were infected with indicated viruses at a MOI of 0.001 in serum free DMEM for 1 h at 37C. After 1 h, virus was removed and virus overlay media (serum free DMEM, 0.14% fraction V BSA, 1x Penicillin/Streptomycin, 1ug/ml TPCK-treated trypsin) containing the same concentration of Universal IFN was added for continuous immune priming of cells. Virus in supernatant of infected cells at indicated timepoints was titrated by plaque assay on MDCK cells.

### ISRE-luciferase reporter assays

The ISRE-luciferase bovine udder fibroblasts were infected with r-EA-2000-C, r-EA-2021-AB, r-Avian-B1.1, r-BHG/NLD/17, r-Goose-B3.13, r-EA-2022-BB, r-Avian-D1.1, r-Bovine-B3.13 and r-Bovine-D1.1 at an MOI of 5 PFU/cell by spinoculation at 500 × *g* for 1h at 12°C. 24 hpi cells were lysed in 200 μl of 1X-passive lysis buffer (Dual-Luciferase reporter assay system, Promega). Post freeze-thawing, readings for firefly luciferase were obtained as mentioned previously for the polymerase assays.

### Protein shutoff assays

To examine protein shutoff, nascent proteins were labelled with puromycin as previously described^122^. Bovine skin fibroblasts were infected with r-Bovine-B3.13, r-Tx/37-B3.13, r-Goose-B3.13, r-EA-2020-C and r-EA-2022-B at a MOI of 5 PFU/cell by spinoculation at 500 × *g* for 1h at 12°C. 24 hpi, puromycin was added to the cells at a final concentration of 3.3 ng/mL for 1 h. Following cell lysis, samples were subjected to SDS-PAGE, and immunoblotting was performed for puromycin, viral PB2, NP & NS1 and β-actin. Quantification of puromycin was performed by normalizing against levels of β-actin.

### Mouse infections

All mice were purchased from Charles River Laboratories (United Kingdom) and housed at the CRUK Scotland Institute. Mice (male, 8-week-old C57BL/6) were intranasally infected with 100 PFU of virus and monitored daily. At post-mortem, lungs were perfused and inflated with agarose and subsequently fixed in 8 % formalin for 24 hours.

### Immunohistochemistry

Formalin-fixed and paraffin embedded (FFPE) mice lung and brain tissue blocks, or precision cut udder slices were cut into ∼3 μm thick sections and mounted on glass slides for immunohistochemistry (IHC) as described previously^123^. The antibodies used were anti-Influenza A H1N1 Nucleoprotein (NP; A/WSN/1933; Novus Biologicals), anti-Cluster of Differentiation 3 (CD3; Agilent Dako), anti-Paired box protein-5 (Pax-5; Abcam). To detect Mx-1, a custom-made monoclonal anti-Mx1 antibody was used (kindly provided by Georg Kochs and Hartmut Hengel, Universitaetsklinikum Freiburg, Germany). As a negative control, the primary antibody was replaced by Dako Real Antibody Diluent (Agilent Dako; Code S2022). Visualisation was performed using the Dako EnVision+ System-HRP Labelled Polymer Anti-Rabbit (Agilent Dako; code K4001) in an automated stainer (Dako Autostainer Link 48, Agilent Technologies). 3,3’-Diaminobenzidine (DAB) was used as a chromogen in all IHC experiments. For confocal imaging, the same primary antibodies (with the same concentrations) have been used as above. Secondary antibodies included AlexaFluor-488 and −594 (codes A11034, A21203; Thermo Fisher Scientific). Dapi was used for nuclear staining. Images have been captured with a Zeiss LSM 710 Confocal Microscope (Zeiss, Wetzlar, Germany).

### Digital Pathology

Slides were digitised and scanned at 20x magnification using the Aperio Versa 8 Slidescanner and Aperio Versa 1.0.4.125 software (Leica Biosystems) for subsequent image analysis. Images of digitised slides were captured using the Aperio ImageScope software v12.4.3.5008 (Leica Biosystems). Positive cell detection and quantification of CD3 and Pax-5, as well as positive pixel quantification of NP, were performed using QuPath software (version 0.4.3;^124^) with settings adjusted to optimise signal detection for each marker^123^. To create the thresholder, the ‘Resolution’ was set to 1.09 μm/pixel or higher. For the ‘Channel’, we used ‘DAB’ (IHC); the ‘Prefilter’ was always ‘Gaussian’ while ‘Smoothing sigma’ as well as the ‘Threshold’ have been tuned for each set of experiments to optimise the correct detection. The ‘Above threshold’ option was always set to ‘Positive’ whereas the ‘Below threshold’ was always set to ‘Negative’. The ‘Region’ to be analysed was set to ‘Any annotation ROI’. The readout is a percentage of positive pixels per total pixels in the annotated area.

The ‘Positive cell’ detection feature has been used instead for the detection of cell membrane-associated CD3-positive cells and nucleus-localised Pax-5. The settings for the ‘Requested pixel size’ were adjusted to 0.5 μm or smaller and the ‘Nucleus parameters’ included a ‘Background radius’ of 8 μm. The ‘Median filter radius’ was set to 0.8 μm or smaller, ‘Sigma’ to 1.5 μm or smaller while the ‘Minimum’ and ‘Maximum’ areas were adjusted to 3–5 μm2 and 70-200 μm2 respectively. The ‘Intensity parameters’ included a ‘Threshold’ of 0.1 and a ‘Max. background intensity’ of 2. The ‘Cell parameters’ included a ‘Cell expansion’ of 3-4.4286 μm. The readout is a percentage of positive cells detected per all cells in the annotated area.

### Statistical analysis

Statistical analyses were performed using GraphPad Prism 10. Data were assessed for Gaussian distribution using the Shapiro-Wilk test or were Log10 transformed. Data from replication kinetics experiments were compared at each time point. Statistical significance was assessed as described in the figure legends. In all analyses statistical significance was determined to be P < 0.05.

## Acknowledgements

We gratefully acknowledge all data contributors, i.e., the authors and their originating laboratories responsible for obtaining the specimens, and their submitting laboratories for generating the genetic sequence and metadata and sharing via the GISAID Initiative, on which this research is based. We thank Lynn Marion Stevenson, Lynn Oxford, Frazer Bell and Lewis Kidd from the Veterinary Diagnostic Service of the University of Glasgow for the excellent support with the tissue sample preparations and staining. We thank Wendy Barclay, Paul Digard and members of the Flu:TrailMap-One Health consortium for insightful discussions. We acknowledge funding from MRC/UKRI (Grant Ref: MR/Y03368X/1) to the to the CVR (MC_UU_00034/3, MC_UU_00034/5, MC_UU_00034/9; to M.P, P.R.M, A.D.F.), and to the G2P2 consortium (MR/Y004205), and the Wellcome Trust to the G2P-global consortium (226141/Z/22/Z). We acknowledge the Flu:TrailMap-One Health consortium, BBSRC FLU-Trailmap (grant Ref: BB/Y007298/1) and Pirbright ISPG (grant Ref: BBS/E/PI/230001B) (to M.P, P.R.M and M.I). Additional funding was supported by CRUK core funding to the Beatson Institute (A31287), CRUK core funding to E.W.-R. (A1920), and the DURABLE consortium (EU4Health Programme 101102733) to R.A.M.F.

---

**Fig S1.**
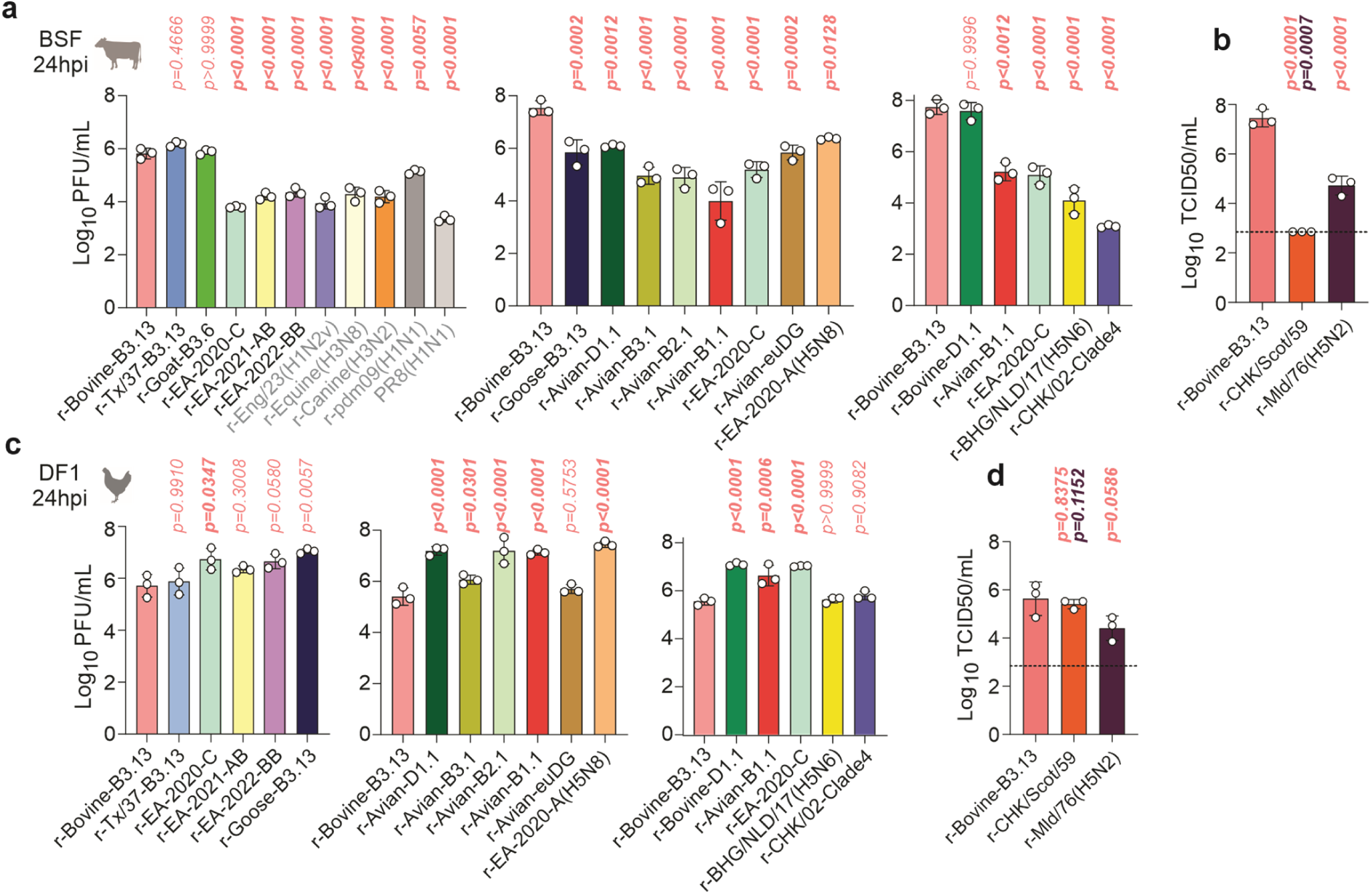
Replication of 2.3.4.4b and other IAV reassortant viruses in Bovine skin fibroblasts (BSF). **a**, BSF were infected with 0.001 PFU per cell and infectious virus in the supernatant at 24 hours post infection (hpi) was quantified by plaque assay on MDCK cells, or by TCID50 on DF-1 cells **b**. **c-d**, Same as above (a-b), but chicken embryonic fibroblast (DF1) cells were infected instead. Data are mean +/- s.d. of three independent biological experiments (n = 3). Data were log-transformed and confirmed to be normally distributed using the Shapiro-Wilk Test. Multiple comparisons between all reassortants were performed using an ordinary one-way ANOVA with Tukey’s multiple comparison test (α = 0.05), although only comparisons relative to controls are shown. P-values in bold indicate statistical significance. H5N1 viruses are shown in black, while labels for non-H5N1 IAVs are shown in grey.

**Fig S2.**
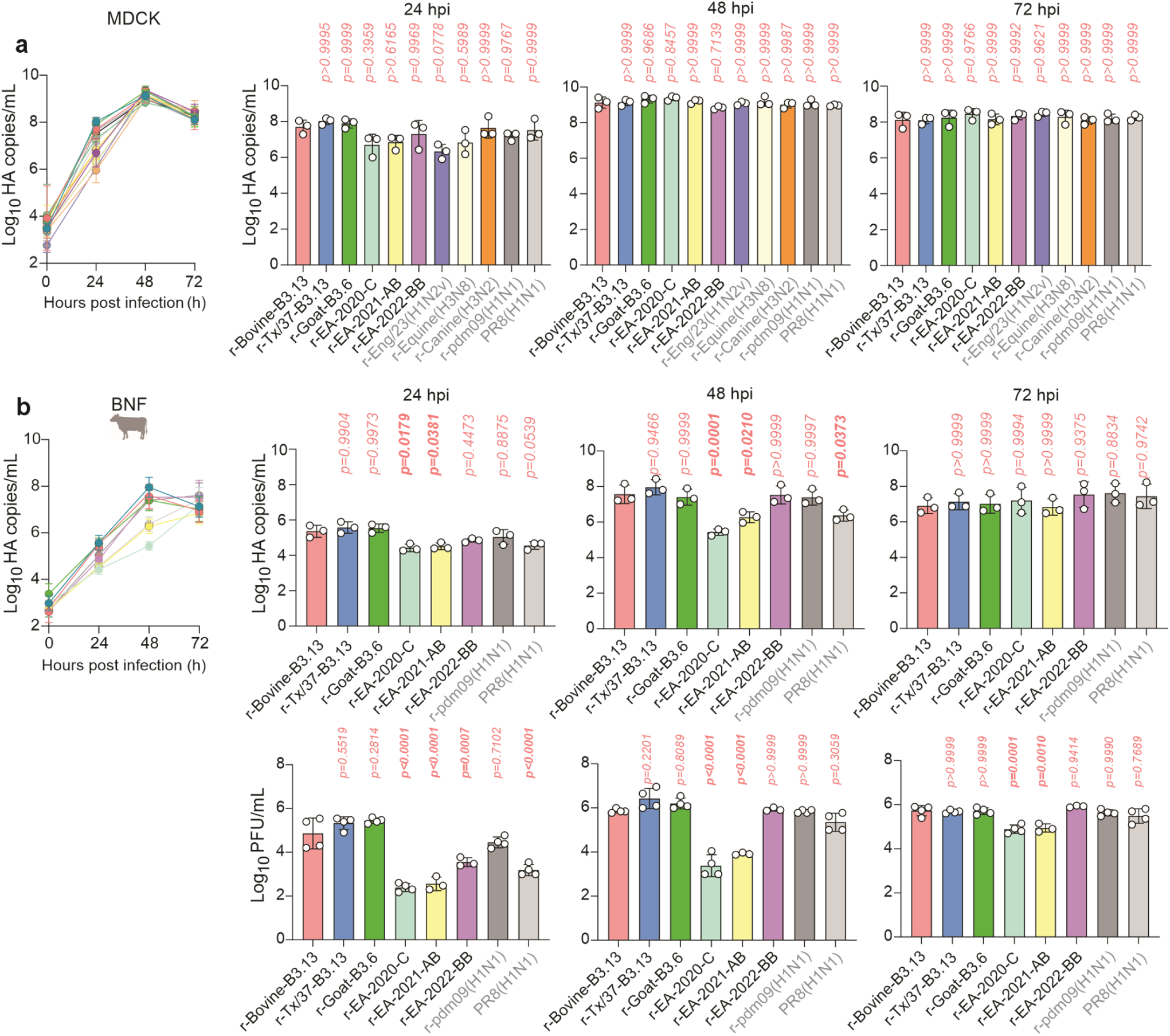
Replication of 2.3.4.4b and other IAV reassortant viruses. **a**, MDCK cells were infected with 0.0005 genome copies per cell and genomic copies in the supernatant at indicated times post infection were titrated by RT-qPCR. **b**, Bovine nasal fibroblasts (BNF) cells were infected as in a and viral genome copies or infectious virus in the supernatant were titrated by RT-qPCR or plaque assay on MDCK cells, respectively. Data are mean +/- s.d. of three independent biological experiments (n = 3). Individual points represent the mean value of two technical duplicates for each biological repeat. Data were log-transformed and confirmed to be normally distributed using the Shapiro-Wilk Test. Multiple comparisons between all groups were performed using an ordinary one-way ANOVA with Tukey’s multiple comparison test (α = 0.05), although only comparisons relative to controls are shown. P-values in bold indicate statistical significance. H5N1 viruses are shown in black, while labels for non-H5N1 IAVs are shown in grey.

**Fig S3.**
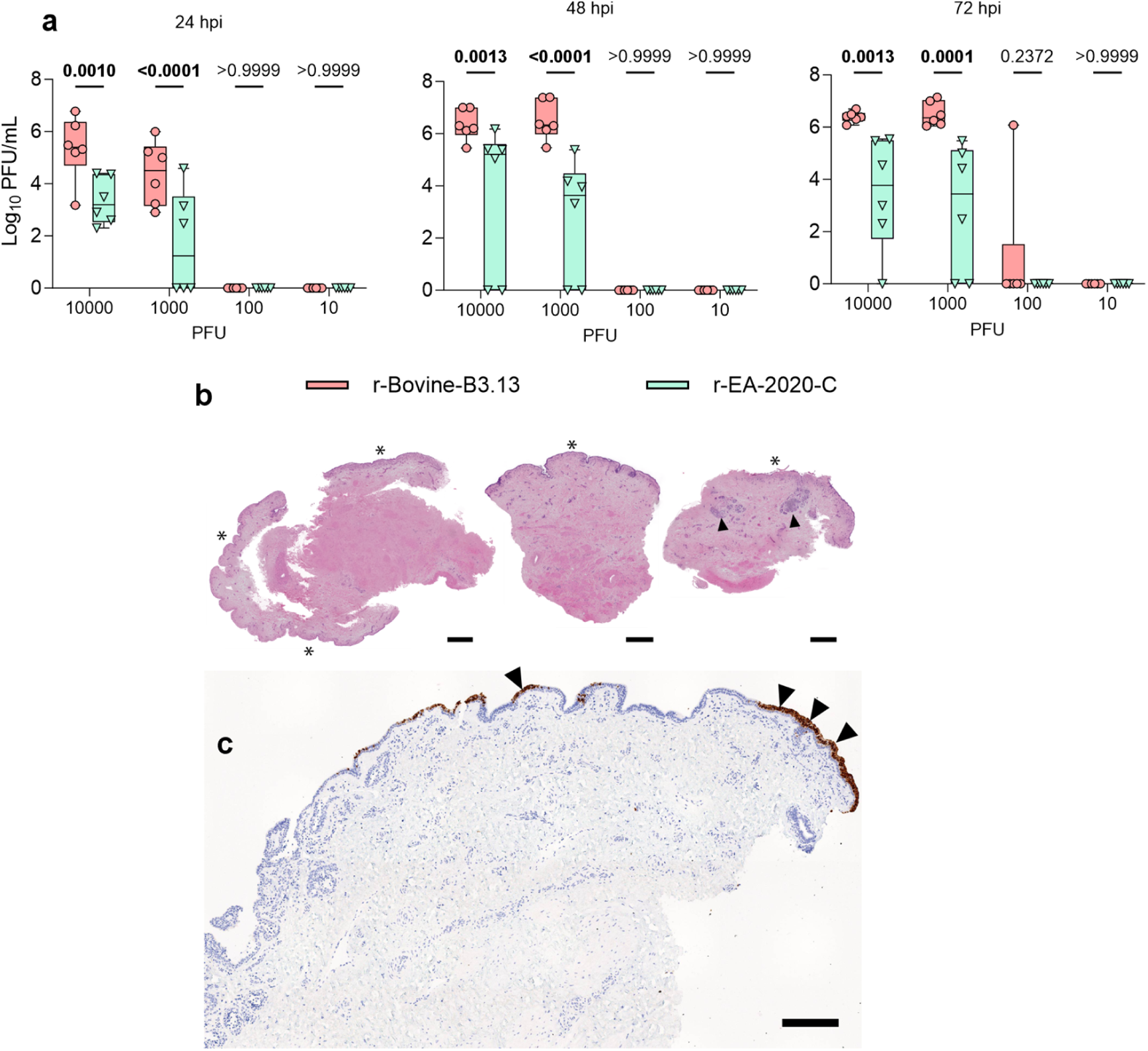
Optimization of Infection of Bovine udder tissue explants. **a**, Udder tissue explants were infected with 10, 100, 1000 or 10000 PFU per explant and infected for indicated time points. Infectious virus in the supernatant was quantified by plaque assay on MDCK cells. Data are from two independently infected tissue explants from three biological donors (n=6). Data were log-transformed and statistical significance between virus groups was determined using a two-way ANOVA with Tukey’s multiple comparison. Boxes represent the interquartile range, the lines indicate the median, and whiskers show the full data range. P-values in bold indicate significance. **b**, Samples show high variability in size and shape: larger sample with larger areas of epithelium surrounding the sample (*, left, infected with r-EA-2022-BB); smaller samples with epithelium on the top only (*, middle, infected with r-Bovine-B3.13); and samples with epithelium on the top (*) and subepithelial glands (arrow heads, right, infected with r-EA-2020-C). **c**, Immunohistochemistry of explants detecting nucleoprotein of influenza virus shows segments of the epithelium being strongly positive in the cytoplasm (arrow heads) next to epithelial areas which are negative. HE staining in **b**, bars in **b** =500 µm. Bar in **c**, 200 µm.

**Fig S4.**
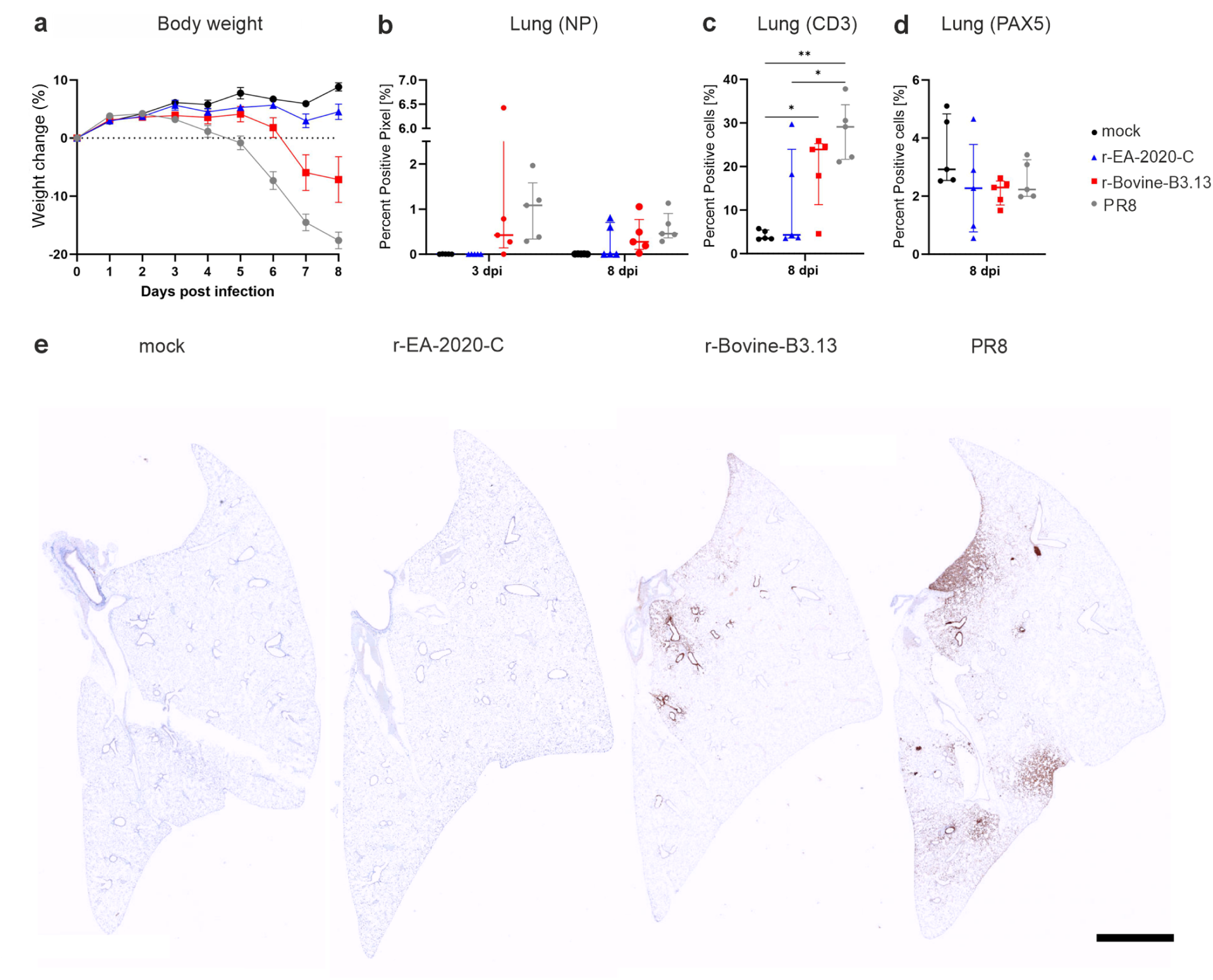
Virulence of 2.3.4.4b reassortant viruses in an experimental mouse model. Groups of 10 C57BL/6 mice were infected with 100 PFU of each recombinant virus, or mock-infected with PBS. 5 mice were euthanized at day 3 and the remaining 5 mice at day 8. **a**, Weights were measured daily. **b** to **d**, Within the lung the amount of viral protein (NP, **b**), the numbers of B cells (CD3, **c**) or T cells (Pax-5, **d**), respectively, was quantified in whole scanned slides of FFPE lung sections and calculated as positive pixel (NP, **b**) or positive cells (CD3, **c**; Pax-5, **d**) per lung area. **e**, Immunohistochemistry of lung sections collected from mock-infected and infected mice, r-EA-2020-C, r-Bovine-B3.13 and PR8, respectively, showing virus positive cells (NP) with brown signal in the bronchi and in the parenchyma (bar, 200 µm). In **b**) Statistical significance was determined using a repeated measures two-way ANOVA with Tukey’s multiple comparison test (α= 0.05). Data in **c**) and **d**) were confirmed to be normally distributed using the Shapiro-Wilk Test and an ordinary one-way ANOVA with Tukey’s multiple comparison test (α = 0.05) was used to compare groups

**Fig S5.**
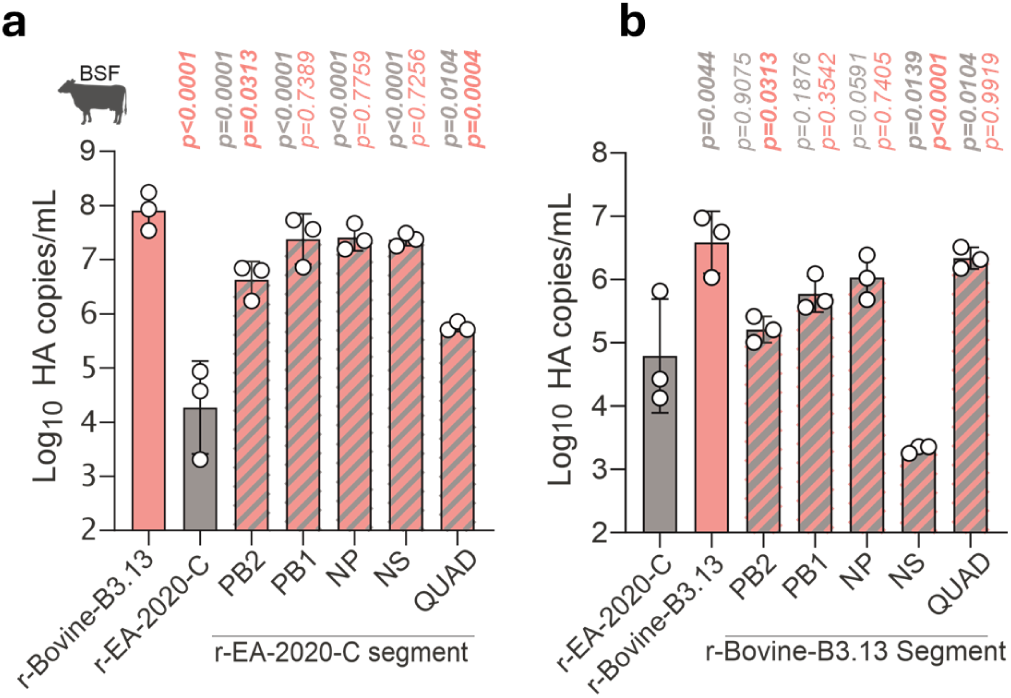
Contribution of B3.13 internal genes to virus replication in bovine cells. **a & b**, BSF cells were infected with 2:6 viruses bearing internal gene reassortant constellations between the ancestral r-EA-2020-C virus and r-Bovine-B3.13. The PB2, PB1, NP or NS segments, or all four (‘QUAD’), were swapped reciprocally between the two. Cells were infected at an MOI of 0.0005 genome copies/cell and viral load in the supernatant at 24h post infection was determined by RT-qPCR as genome copies/mL. Data are mean +/- s.d. of three independent experiments. Each dot represents the mean of two technical duplicates from the three biological repeats. Data were log-transformed and confirmed to be normally distributed using the Shapiro-Wilk Test. Multiple comparisons between all viruses were performed using an ordinary one-way ANOVA with Tukey’s multiple comparison test (α = 0.05). Only comparisons relative to controls are shown. P-values in bold indicate statistical significance.

**Figure S6.**
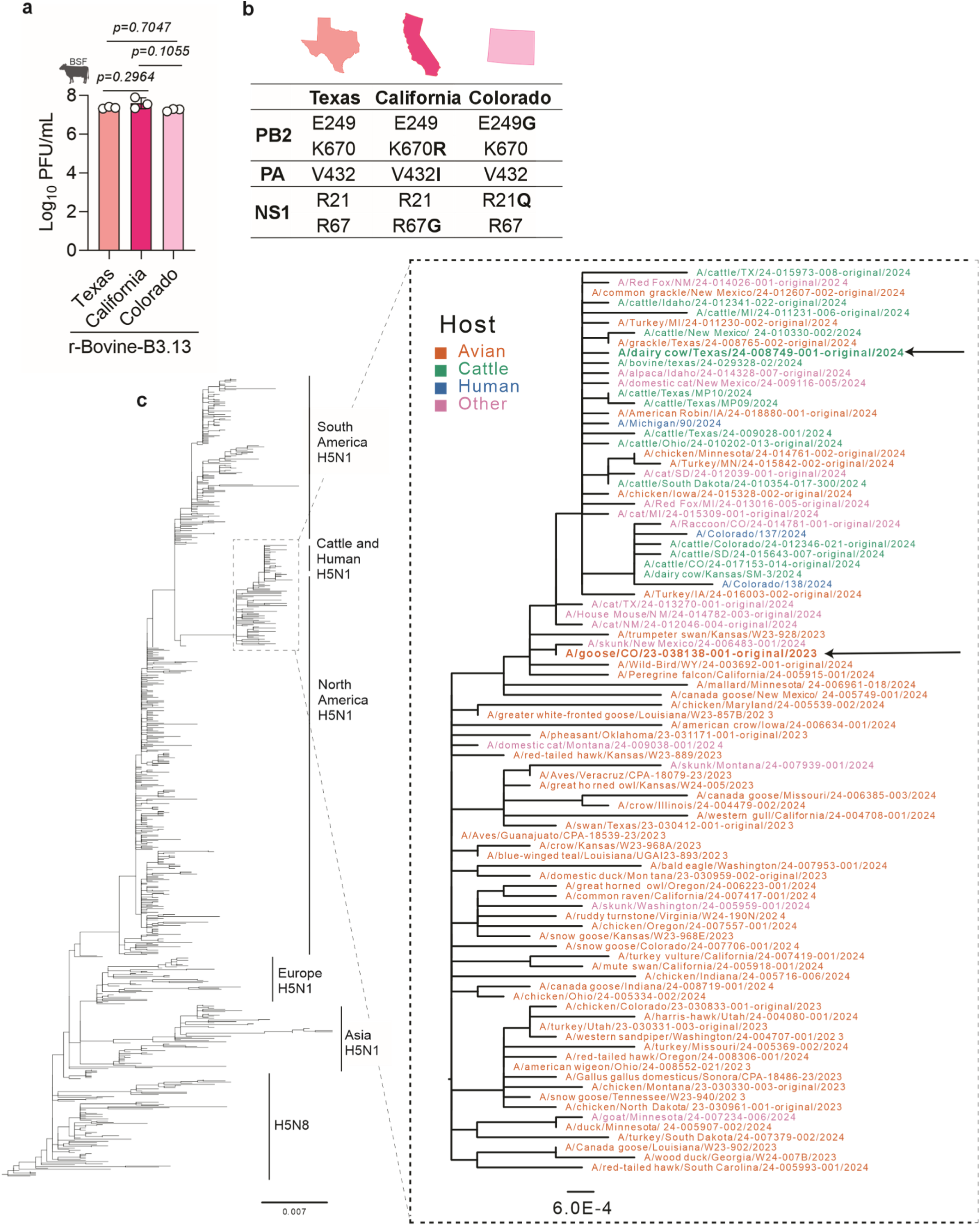
Evolution and Replication of r-Bovine-B3.13 viruses that evolved during the outbreak. **a**, Replication of r-Bovine-B3.13 Texas, Colorado and California virus strains in BSF cells at 24 hpi. Cells were infected with an MOI of 0.001 PFU/mL and virus titres in the supernatant were determined by plaque assay on MDCK cells. Data were log-transformed and assessed for normality using the Shapiro-Wilk Test. Statistical comparisons between all viruses were performed using an ordinary one-way ANOVA with Tukey’s multiple comparison test (α = 0.05). P values in bold indicate significance. **b**, Table summarizing the amino acid substitutions acquired during the evolution of Bovine-B3.13 in dairy cattle throughout the outbreak. **c**, Maximum likelihood phylogeny of H5N1 HA de-duplicated amino acid sequences focusing on the evolution of the European and American 2.3.4.4b lineages. The part of the tree associated to the outbreak in dairy cattle is expanded and the tips are coloured according to the host. The tree was reconstructed using IQ-TREE2 with the best model selected by BIC (Bayesian Information Criterion).

**Fig S7.**
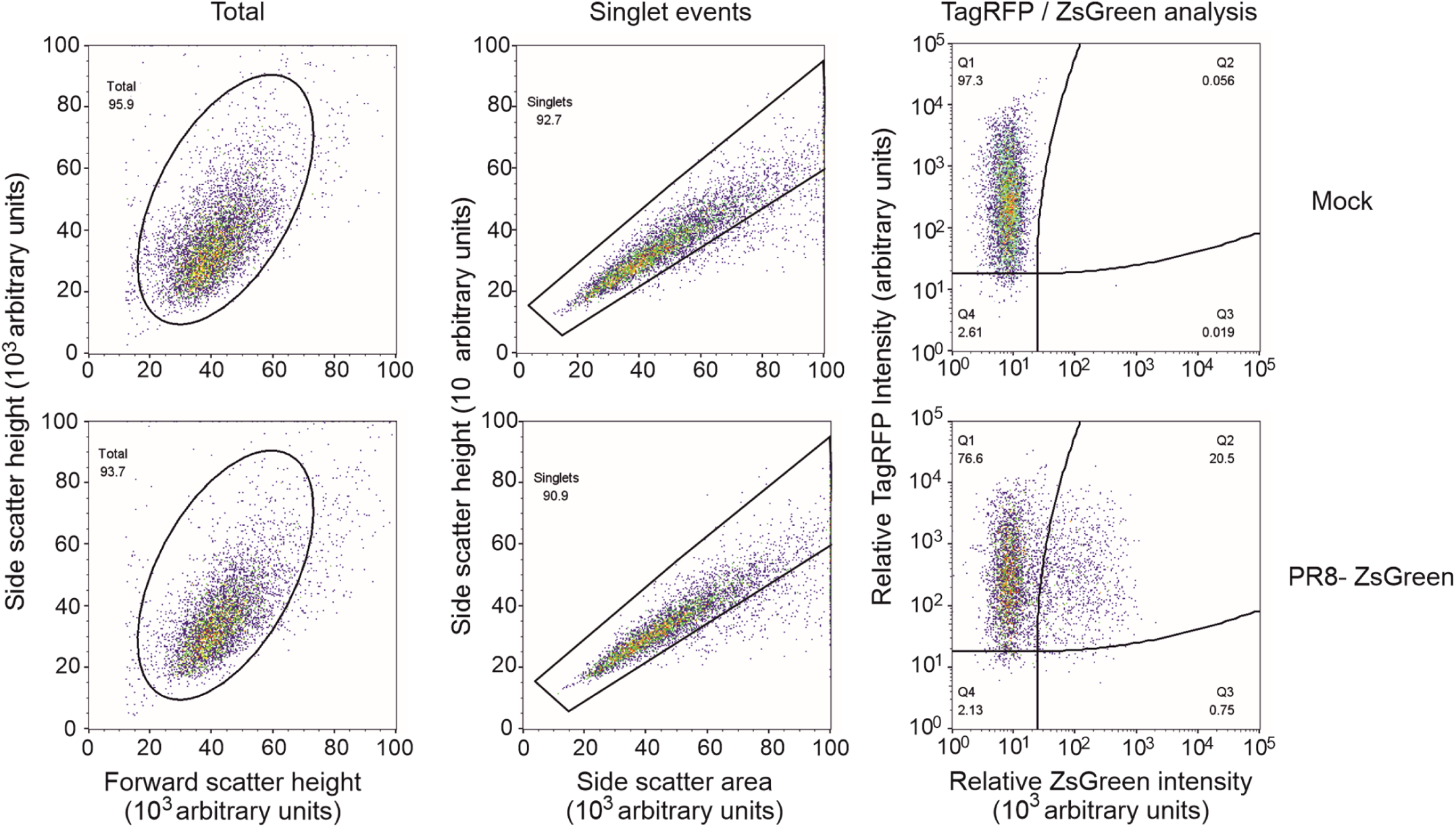
Example flow cytometry gating of an IAV-ZsGreen titration on A549-SCRPSY-Mx1 cells. Population gating was performed on FlowJo. Shown is an example of A549-Hs. MxA expressing cells either mock-infected (top row) or infected with PR8-ZsGreen (bottom row). A ‘total’ cell population (leftmost plots) was initially gated using forward and side scatter height intensity with a threshold applied to the forward scatter height to remove false events from debris or other particles. Single cells from this population (middle plots) were gated using side scatter height and area. Relative TagRFP (expressed by the SCRPSY-Mx1 lentiviral vector) and ZsGreen (expressed following IAV-ZsGreen virus infection) intensities were measured from this single cell population. A mock-infected sample was used to set the ZsGreen thresholding to gate for ZsGreen-positive events. The Q2 and Q3 plots were summed to calculate IAV infection percentage. The numerical values in each quadrant of each plot represents the percentage of cells in that population.

**Fig S8.**
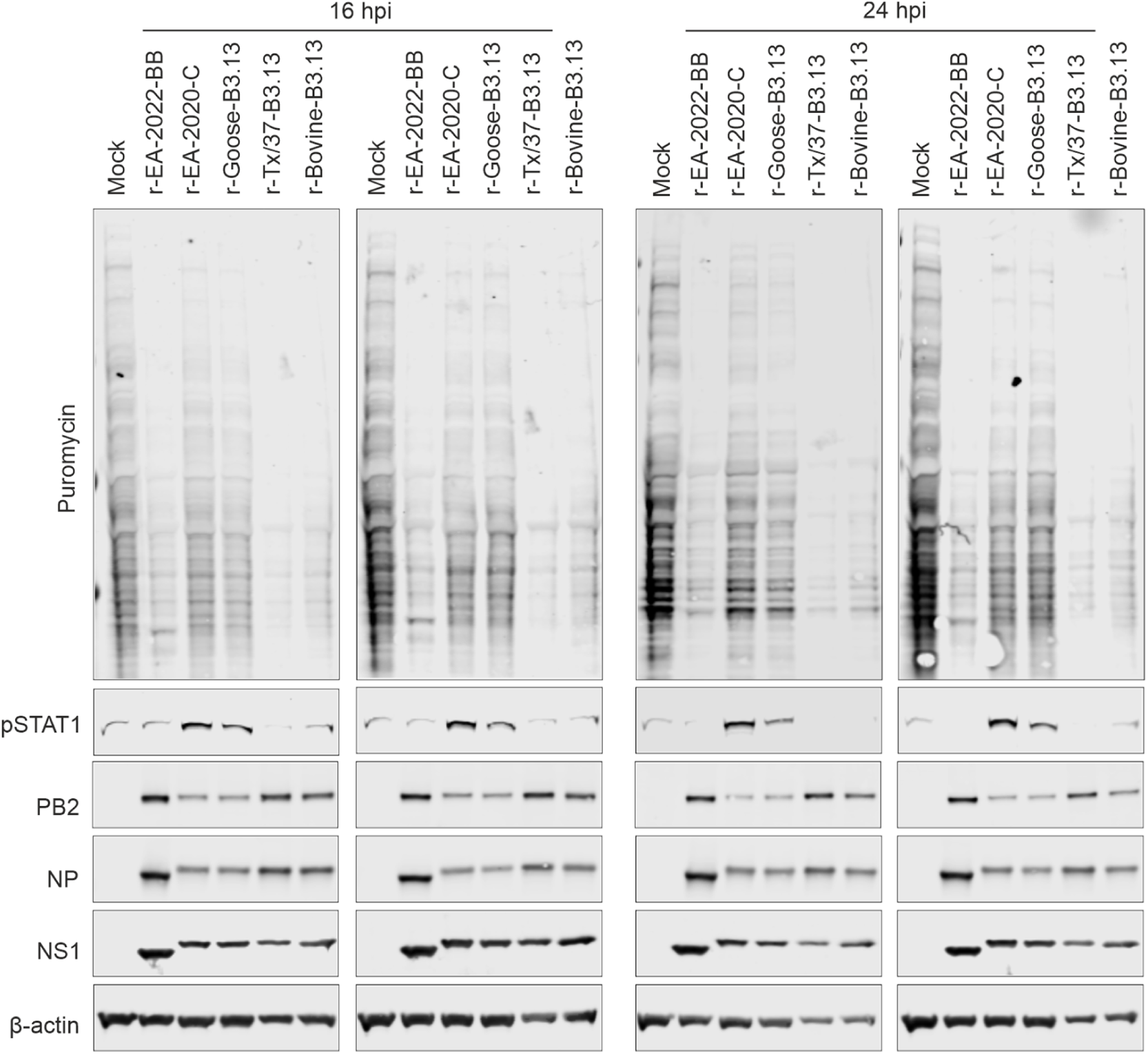
Host-shutoff activity of 2.3.4.4b reassortant viruses in bovine cells. Immortalised BSF were infected at an MOI of 5 PFU/cell for 16 h or 24 h and pulsed with puromycin for 1 hour. Cell lysates were assessed by western blot for total cellular proteins, phosphorylated STAT1 and the viral proteins PB2, NP and NS1 following puromycin-labelling. β-actin was assessed as a loading control.

**Table S1.**
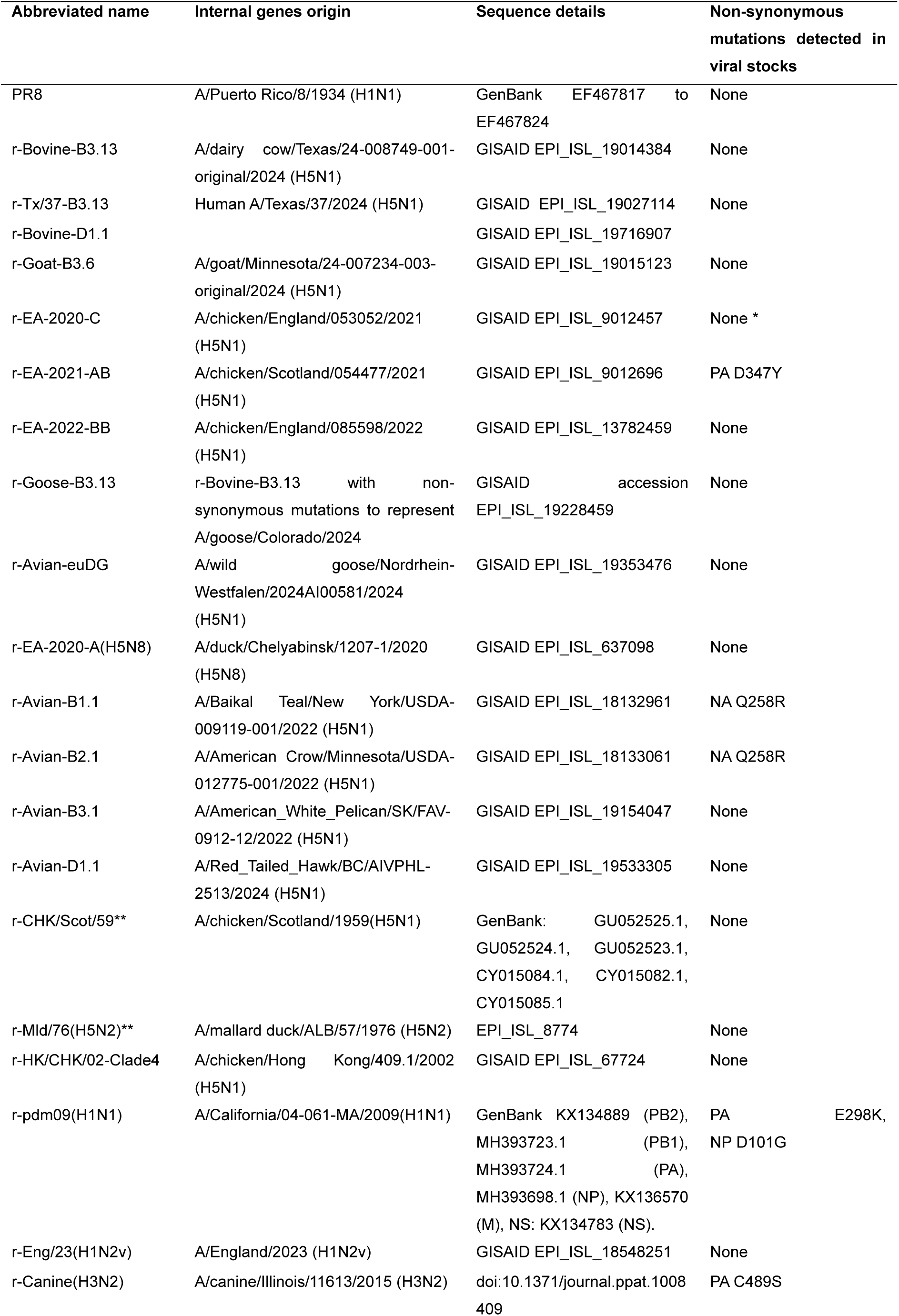
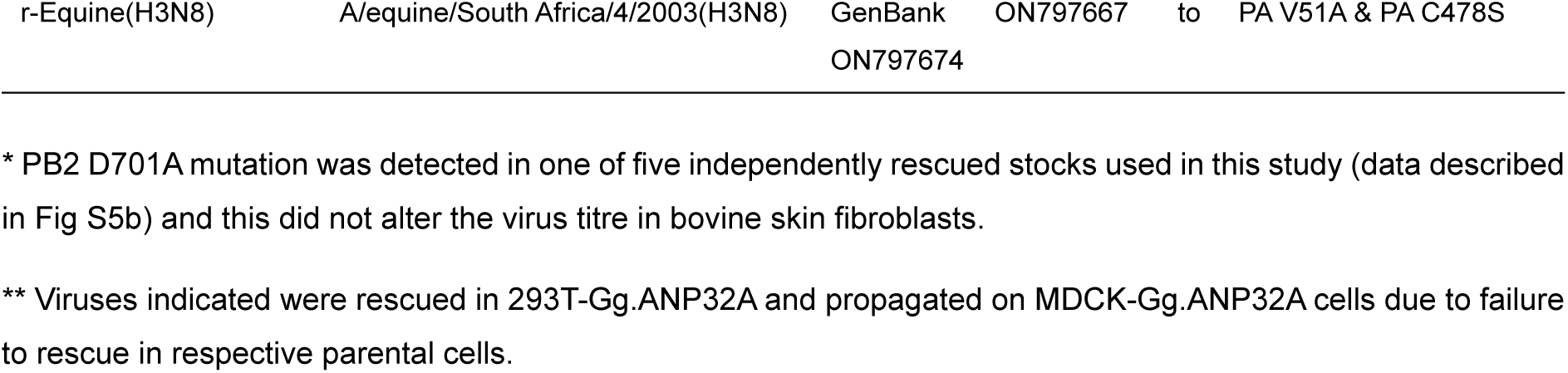
2:6 Reassortants used in this study. Each virus has the HA and NA segments from PR8, and all remaining segments from the virus listed.

| Abbreviated name | Internal genes origin | Sequence details | Non-synonymous mutations detected in viral stocks |
| --- | --- | --- | --- |
| PR8 | A/Puerto Rico/8/1934 (H1N1) | GenBank EF467817 to EF467824 | None |
| r-Bovine-B3.13 | A/dairy cow/Texas/24-008749-001-original/2024 (H5N1) | GISAID EPI_ISL_19014384 | None |
| r-Tx/37-B3.13 | Human A/Texas/37/2024 (H5N1) | GISAID EPI_ISL_19027114 | None |
| r-Bovine-D1.1 |  | GISAID EPI_ISL_19716907 |  |
| r-Goat-B3.6 | A/goat/Minnesota/24-007234-003-original/2024 (H5N1) | GISAID EPI_ISL_19015123 | None |
| r-EA-2020-C | A/chicken/England/053052/2021 (H5N1) | GISAID EPI_ISL_9012457 | None * |
| r-EA-2021-AB | A/chicken/Scotland/054477/2021 (H5N1) | GISAID EPI_ISL_9012696 | PA D347Y |
| r-EA-2022-BB | A/chicken/England/085598/2022 (H5N1) | GISAID EPI_ISL_13782459 | None |
| r-Goose-B3.13 | r-Bovine-B3.13 with non-synonymous mutations to represent A/goose/Colorado/2024 | GISAID accession EPI_ISL_19228459 | None |
| r-Avian-euDG | A/wild goose/Nordrhein-Westfalen/2024AI00581/2024 (H5N1) | GISAID EPI_ISL_19353476 | None |
| r-EA-2020-A(H5N8) | A/duck/Chelyabinsk/1207-1/2020 (H5N8) | GISAID EPI_ISL_637098 | None |
| r-Avian-B1.1 | A/Baikal Teal/New York/USDA-009119-001/2022 (H5N1) | GISAID EPI_ISL_18132961 | NA Q258R |
| r-Avian-B2.1 | A/American Crow/Minnesota/USDA-012775-001/2022 (H5N1) | GISAID EPI_ISL_18133061 | NA Q258R |
| r-Avian-B3.1 | A/American_White_Pelican/SK/FAV-0912-12/2022 (H5N1) | GISAID EPI_ISL_19154047 | None |
| r-Avian-D1.1 | A/Red_Tailed_Hawk/BC/AIVPHL-2513/2024 (H5N1) | GISAID EPI_ISL_19533305 | None |
| r-CHK/Scot/59** | A/chicken/Scotland/1959(H5N1) | GenBank: GU052525.1, GU052524.1, GU052523.1, CY015084.1, CY015082.1, CY015085.1 | None |
| r-Mld/76(H5N2)** | A/mallard duck/ALB/57/1976 (H5N2) | EPI_ISL_8774 | None |
| r-HK/CHK/02-Clade4 | A/chicken/Hong Kong/409.1/2002 (H5N1) | GISAID EPI_ISL_67724 | None |
| r-pdm09(H1N1) | A/California/04-061-MA/2009(H1N1) | GenBank KX134889 (PB2), MH393723.1 (PB1), MH393724.1 (PA), MH393698.1 (NP), KX136570 (M), NS: KX134783 (NS). | PA E298K, NP D101G |
| r-Eng/23(H1N2v) | A/England/2023 (H1N2v) | GISAID EPI_ISL_18548251 | None |
| r-Canine(H3N2) | A/canine/Illinois/11613/2015 (H3N2) | doi:10.1371/journal.ppat.1008409 | PA C489S |

\* PB2 D701A mutation was detected in one of five independently rescued stocks used in this study (data described in Fig S5b) and this did not alter the virus titre in bovine skin fibroblasts.
\*\* Viruses indicated were rescued in 293T-Gg.ANP32A and propagated on MDCK-Gg.ANP32A cells due to failure to rescue in respective parental cells.

**Table S2.** 4:4 Reassortants used in this study. Each virus has the HA, NA, M and NS segments from PR8, and all remaining segments from the virus listed.

| Abbreviated name | Internal genes origin | Sequence details | Non-synonymous mutations detected in viral stocks |
| --- | --- | --- | --- |
| r-Mld/99(H1N1) | A/mallard/Netherlands/10-Cam/1999(H1N1) | GenBank KC209512 to KC209519 | PA C489S |
| r-Bovine-B3.13* | A/dairy cow/Texas/24-008749-001-original/2024 (H5N1) | GISAID EPI_ISL_19014384 | None |
| r-Tx/37-B3.13* | Human A/Texas/37/2024 (H5N1) | GISAID EPI_ISL_19027114 | None |
| r-Goat-B3.6* | A/goat/Minnesota/24-007234-003-original/2024 (H5N1) | GISAID EPI_ISL_19015123 |  |
| r-EA-2020-C* | A/chicken/England/053052/2021 (H5N1) | GISAID EPI_ISL_9012457 |  |
| r-EA-2021-AB* | A/chicken/Scotland/054477/2021 (H5N1) | GISAID EPI_ISL_9012696 | None |
| r-EA-2022-BB* | A/chicken/England/085598/2022 (H5N1) | GISAID EPI_ISL_13782459 | None |
| r-Goose-B3.13 | r-Bovine-B3.13 with non-synonymous mutations to represent A/goose/Colorado/2024 | GISAID accession EPI_ISL_19228459 |  |
| r-Eng/23(H1N2v) | A/England/2023 (H1N2v) | GISAID EPI_ISL_18548251 | None |
| r-Anhui/13(H7N9) | A/Anhui/2013 (H7N9) | GISAID EPI_ISL_17264823 | None |
| r-Iowa/11(H3N2v) | A/Iowa/08/2011 | EPI_ISL_99214 | PB1 D638V |
| r-Ohio/16(H3N2v) | A/Ohio/27/2016 | OH/16: EPI_ISL_232044 | None |
| r-Chn/23(H3N8) | A/Guangdong/ZS-2023SF005/2023 | EPI_ISL_17464053 | None |
| r-Arg(H4N2) | A/silver teal/Argentina/CIP051-32/2011 | OQ821651.1, OQ821652.1, OQ821653.1, OQ821655.1 | None |
| r-Nld(H7N7) | A/Netherlands/219/03(H7N7) | AY342413.1, AY340083.1, AY342418.1, AY342425.1 | None |

\*Viruses indicated were rescued in 293T-Gg.ANP32A and propagated on MDCK-Gg.ANP32A cells due to failure to rescue in respective parental cells without incurring known mammalian-adaptive mutations

**Table S3.**
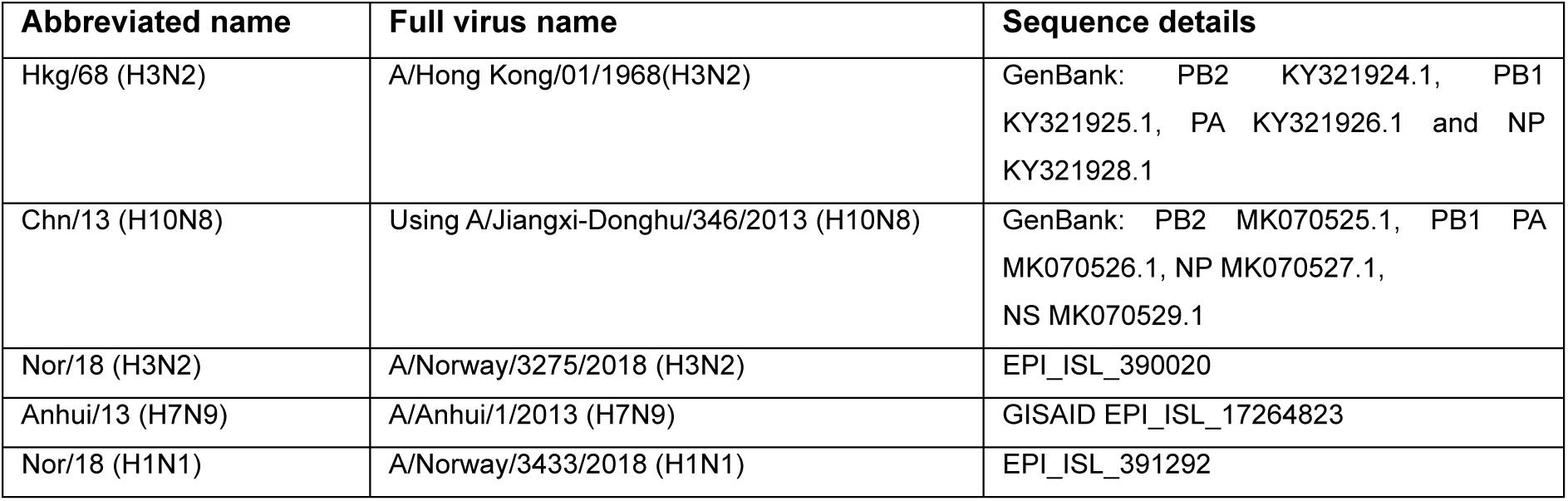
Viruses tested exclusively in polymerase activity assays in this study. The polymerase of these viruses was tested in polymerase activity assays described in Fig 3.

| Abbreviated name | Full virus name | Sequence details |
| --- | --- | --- |
| Hkg/68 (H3N2) | A/Hong Kong/01/1968(H3N2) | GenBank: PB2 KY321924.1, PB1 KY321925.1, PA KY321926.1 and NP KY321928.1 |
| Chn/13 (H10N8) | Using A/Jiangxi-Donghu/346/2013 (H10N8) | GenBank: PB2 MK070525.1, PB1 PA MK070526.1, NP MK070527.1, NS MK070529.1 |
| Nor/18 (H3N2) | A/Norway/3275/2018 (H3N2) | EPI_ISL_390020 |
| Anhui/13 (H7N9) | A/Anhui/1/2013 (H7N9) | GISAID EPI_ISL_17264823 |
| Nor/18 (H1N1) | A/Norway/3433/2018 (H1N1) | EPI_ISL_391292 |

## References

1 Webster, R. G., Bean, W. J., Gorman, O. T., Chambers, T. M. & Kawaoka, Y. Evolution and ecology of influenza A viruses. Microbiol Rev 56, 152–179 (1992).

2 Worobey, M., Han, G. Z. & Rambaut, A. Genesis and pathogenesis of the 1918 pandemic H1N1 influenza A virus. Proc Natl Acad Sci U S A 111, 8107–8112 (2014). 10.1073/pnas.1324197111

3 Scholtissek, C., Rohde, W., Von Hoyningen, V. & Rott, R. On the origin of the human influenza virus subtypes H2N2 and H3N2. Virology 87, 13–20 (1978).

4 Kawaoka, Y., Krauss, S. & Webster, R. G. Avian-to-human transmission of the PB1 gene of influenza A viruses in the 1957 and 1968 pandemics. J Virol 63, 4603–4608 (1989). 10.1128/jvi.63.11.4603-4608.1989

5 Taubenberger, J. K. et al. Characterization of the 1918 influenza virus polymerase genes. Nature 437, 889–893 (2005). http://www.nature.com/nature/journal/v437/n7060/suppinfo/nature04230_S1.html

6 Patrono, L. V. et al. Archival influenza virus genomes from Europe reveal genomic variability during the 1918 pandemic. Nat Commun 13, 2314 (2022). 10.1038/s41467-022-29614-9

7 Peacock, T. et al. The global H5N1 influenza panzootic in mammals. Nature (2024). 10.1038/s41586-024-08054-z

8 Pohlmann, A. et al. Has epizootic become enzootic? Evidence for a fundamental change in the infection dynamics of highly pathogenic avian influenza in Europe, 2021. MBio 13, e00609-00622 (2022).

9 Caliendo, V. et al. Transatlantic spread of highly pathogenic avian influenza H5N1 by wild birds from Europe to North America in 2021. Scientific reports 12, 11729 (2022).

10 Elsmo, E. J. et al. Highly Pathogenic Avian Influenza A(H5N1) Virus Clade 2.3.4.4b Infections in Wild Terrestrial Mammals, United States, 2022. Emerg Infect Dis 29, 2451-2460 (2023). 10.3201/eid2912.230464

11 Jakobek, B. T. et al. Influenza A(H5N1) Virus Infections in 2 Free-Ranging Black Bears (Ursus americanus), Quebec, Canada. Emerg Infect Dis 29, 2145–2149 (2023). 10.3201/eid2910.230548

12 Moreno, A. et al. Asymptomatic infection with clade 2.3.4.4b highly pathogenic avian influenza A(H5N1) in carnivore pets, Italy, April 2023. Euro Surveill 28 (2023). 10.2807/1560-7917.ES.2023.28.35.2300441

13 Rabalski, L. et al. Emergence and potential transmission route of avian influenza A (H5N1) virus in domestic cats in Poland, June 2023. Euro Surveill 28 (2023). 10.2807/1560-7917.ES.2023.28.31.2300390

14 Leguia, M. et al. Highly pathogenic avian influenza A (H5N1) in marine mammals and seabirds in Peru. Nature Communications 14, 5489 (2023). 10.1038/s41467-023-41182-0

15 Pardo-Roa, C. et al. Cross-species transmission and PB2 mammalian adaptations of highly pathogenic avian influenza A/H5N1 viruses in Chile. bioRxiv (2023). 10.1101/2023.06.30.547205

16 Uhart, M. et al. Massive outbreak of Influenza A H5N1 in elephant seals at Península Valdés, Argentina: increased evidence for mammal-to-mammal transmission. bioRxiv, 2024.2005.2031.596774 (2024). 10.1101/2024.05.31.596774

17 Tomás, G. et al. Highly pathogenic avian influenza H5N1 virus infections in pinnipeds and seabirds in Uruguay: Implications for bird-mammal transmission in South America. Virus Evol 10, veae031 (2024). 10.1093/ve/veae031

18 Burrough, E. R. et al. Highly Pathogenic Avian Influenza A(H5N1) Clade 2.3.4.4b Virus Infection in Domestic Dairy Cattle and Cats, United States, 2024. Emerg Infect Dis 30 (2024). 10.3201/eid3007.240508

19 Kalthoff, D., Hoffmann, B., Harder, T., Durban, M. & Beer, M. Experimental infection of cattle with highly pathogenic avian influenza virus (H5N1). Emerg Infect Dis 14, 1132–1134 (2008). 10.3201/eid1407.071468

20 Sreenivasan, C. C., Thomas, M., Kaushik, R. S., Wang, D. & Li, F. Influenza A in Bovine Species: A Narrative Literature Review. Viruses 11 (2019). 10.3390/v11060561

21 Caliendo, V. et al. Transatlantic spread of highly pathogenic avian influenza H5N1 by wild birds from Europe to North America in 2021. Scientific Reports 12, 11729 (2022). 10.1038/s41598-022-13447-z

22 Caserta, L. C. et al. Spillover of highly pathogenic avian influenza H5N1 virus to dairy cattle. Nature 634, 669–676 (2024). 10.1038/s41586-024-07849-4

23 Worobey, M., Gangavarapu K., Pekar, JE., Joy JB., Moncla L., Kraemer MUG., Dudas, G., Goldhill, D., Ruis, C., Serrano, LM., Ji, X., Andersen, KG., Wertheim JO., Lemey, P., Suchard, MA., Rasmussen, AL., Chand, M., Groves, N., Pybus, OG., Peacock, TP., Rambaut, A., Nelson, MI. Preliminary report on genomic epidemiology of the 2024 H5N1 influenza A virus outbreak in U.S. cattle., <https://virological.org/t/preliminary-report-on-genomic-epidemiology-of-the-2024-h5n1-influenza-a-virus-outbreak-in-u-s-cattle-part-1-of-2/970/1> (2024).

24 Mostafa, A., Nogales, A. & Martinez-Sobrido, L. Highly pathogenic avian influenza H5N1 in the United States: recent incursions and spillover to cattle. Npj Viruses 3, 54 (2025). 10.1038/s44298-025-00138-5

25 Nguyen, T. Q. et al. Emergence and interstate spread of highly pathogenic avian influenza A(H5N1) in dairy cattle in the United States. Science 388, eadq0900 (2025). 10.1126/science.adq0900

26 Nguyen, T.-Q. et al. Emergence and interstate spread of highly pathogenic avian influenza A(H5N1) in dairy cattle. bioRxiv, 2024.2005.2001.591751 (2024). 10.1101/2024.05.01.591751

27 CDC. H5 Bird Flu: Current Situation <https://www.cdc.gov/bird-flu/situation-summary/index.html?CDC_AA_refVal=https%3A%2F%2Fwww.cdc.gov%2Fbird-flu%2Fphp%2Favian-flu-summary%2Findex.html https://www.cdc.gov%2Fbird-flu%2Fphp%2Favian-flu-summary%2Findex.html> (2025).

28 APHIS. HPAI Confirmed Cases in Livestock https://www.aphis.usda.gov/livestock-poultry-disease/avian/avian-influenza/hpai-detections/hpai-confirmed-cases-livestock 2025).

29 Worobey, M., Pekar, JE., Peacock, TP., Moncla, L., Gangavarapu, P., Venkatesh, D., Goldhill, DH., Gangavarapu, K., Kraemer, MUG., Dudas, G., Joy, JB., Ruis, C., Xiang, Ji., Chand, M., Groves, N., Pybus, OG., Rasmussen, AL., Wertheim, JO., Rambaut, A., Lemey, P. Timing and molecular characterisation of the transmission to cattle of H5N1 influenza A virus genotype D1.1, clade 2.3.4.4b., <https://virological.org/t/timing-and-molecular-characterisation-of-the-transmission-to-cattle-of-h5n1-influenza-a-virus-genotype-d1-1-clade-2-3-4-4b/991> (2025).

30 APHIS. H5N1 Highly Pathogenic Avian Influenza (HPAI) in Livestock: Information for small ruminant (sheep and goat) and camelid stakeholders <https://www.aphis.usda.gov/sites/default/files/small-ruminant-camelid-h5n1-info.pdf> (2024).

31 Long, J. S., Mistry, B., Haslam, S. M. & Barclay, W. S. Host and viral determinants of influenza A virus species specificity. Nature Reviews Microbiology 17, 67–81 (2019). 10.1038/s41579-018-0115-z

32 Gagneux, P. et al. Human-specific Regulation of α2–6-linked Sialic Acids*. Journal of Biological Chemistry 278, 48245–48250 (2003). 10.1074/jbc.M309813200

33 Shinya, K. et al. Avian flu: influenza virus receptors in the human airway. Nature 440, 435–436 (2006). 10.1038/440435a

34 Matrosovich, M. N., Matrosovich, T. Y., Gray, T., Roberts, N. A. & Klenk, H. D. Human and avian influenza viruses target different cell types in cultures of human airway epithelium. Proc Natl Acad Sci U S A 101, 4620–4624 (2004). 10.1073/pnas.0308001101

35 van Riel, D. et al. H5N1 Virus Attachment to Lower Respiratory Tract. Science 312, 399–399 (2006). 10.1126/science.1125548

36 Fukuyama, S. et al. Multi-spectral fluorescent reporter influenza viruses (Color-flu) as powerful tools for in vivo studies. Nat Commun 6, 6600 (2015). 10.1038/ncomms7600

37 Eisfeld, A. J. et al. Pathogenicity and transmissibility of bovine H5N1 influenza virus. Nature 633, 426–432 (2024). 10.1038/s41586-024-07766-6

38 Kristensen, C., Jensen, H. E., Trebbien, R., Webby, R. J. & Larsen, L. E. Avian and Human Influenza A Virus Receptors in Bovine Mammary Gland. Emerg Infect Dis 30, 1907–1911 (2024). 10.3201/eid3009.240696

39 Ríos Carrasco, M., Gröne, A., van den Brand, J. M. A. & de Vries, R. P. The mammary glands of cows abundantly display receptors for circulating avian H5 viruses. J Virol, e0105224 (2024). 10.1128/jvi.01052-24

40 Nelli, R. K. et al. Sialic Acid Receptor Specificity in Mammary Gland of Dairy Cattle Infected with Highly Pathogenic Avian Influenza A(H5N1) Virus. Emerg Infect Dis 30, 1361–1373 (2024). 10.3201/eid3007.240689

41 Santos, J. J. S. et al. Bovine H5N1 binds poorly to human-type sialic acid receptors. Nature 640, E18–E20 (2025). 10.1038/s41586-025-08821-6

42 Yang, J. et al. The haemagglutinin gene of bovine-origin H5N1 influenza viruses currently retains receptor-binding and pH-fusion characteristics of avian host phenotype. Emerg Microbes Infect 14, 2451052 (2025). 10.1080/22221751.2025.2451052

43 Song, H. et al. Receptor binding, structure, and tissue tropism of cattle-infecting H5N1 avian influenza virus hemagglutinin. Cell 188, 919–929.e919 (2025). 10.1016/j.cell.2025.01.019

44 Lin, T.-H. et al. A single mutation in bovine influenza H5N1 hemagglutinin switches specificity to human receptors. Science 386, 1128–1134 (2024). doi:10.1126/science.adt0180

45 Good, M. R. et al. A single mutation in dairy cow-associated H5N1 viruses increases receptor binding breadth. Nature Communications 15, 10768 (2024). 10.1038/s41467-024-54934-3

46 Chopra, P. et al. Receptor-binding specificity of a bovine influenza A virus. Nature 640, E21–E27 (2025). 10.1038/s41586-025-08822-5

47 Eisfeld, A. J. et al. Pathogenicity and transmissibility of bovine H5N1 influenza virus. Nature (2024). 10.1038/s41586-024-07766-6

48 Russell, C. J., Hu, M. & Okda, F. A. Influenza Hemagglutinin Protein Stability, Activation, and Pandemic Risk. Trends Microbiol 26, 841–853 (2018). 10.1016/j.tim.2018.03.005

49 Blumenkrantz, D., Roberts, K. L., Shelton, H., Lycett, S. & Barclay, W. S. The short stalk length of highly pathogenic avian influenza H5N1 virus neuraminidase limits transmission of pandemic H1N1 virus in ferrets. J Virol 87, 10539–10551 (2013). 10.1128/jvi.00967-13

50 Long, J. S. et al. Species difference in ANP32A underlies influenza A virus polymerase host restriction. Nature 529, 101–104 (2016). 10.1038/nature16474

51 Zhang, H. et al. Fundamental Contribution and Host Range Determination of ANP32A and ANP32B in Influenza A Virus Polymerase Activity. J Virol 93 (13): 10.1128 (2019). 10.1128/jvi.00174-19

52 Long, J. S. et al. Species specific differences in use of ANP32 proteins by influenza A virus. Elife 8:e45066 (2019). 10.7554/eLife.45066

53 Staller, E. et al. ANP32 Proteins Are Essential for Influenza Virus Replication in Human Cells. J Virol 93 (17):10.1128 (2019). 10.1128/jvi.00217-19

54 Zhang, Z. et al. Selective usage of ANP32 proteins by influenza B virus polymerase: Implications in determination of host range. PLOS Pathogens 16, e1008989 (2020). 10.1371/journal.ppat.1008989

55 Carrique, L. et al. Host ANP32A mediates the assembly of the influenza virus replicase. Nature 587, 638–643 (2020). 10.1038/s41586-020-2927-z

56 Gabriel, G. et al. Differential use of importin-α isoforms governs cell tropism and host adaptation of influenza virus. Nature Communications 2, 156 (2011). 10.1038/ncomms1158

57 Pinto, R. M. et al. BTN3A3 evasion promotes the zoonotic potential of influenza A viruses. Nature 619, 338–347 (2023). 10.1038/s41586-023-06261-8

58 Dittmann, J. et al. Influenza A virus strains differ in sensitivity to the antiviral action of Mx-GTPase. J Virol 82, 3624–3631 (2008). 10.1128/jvi.01753-07

59 Zimmermann, P., Manz, B., Haller, O., Schwemmle, M. & Kochs, G. The viral nucleoprotein determines Mx sensitivity of influenza A viruses. J Virol 85, 8133–8140 (2011). 10.1128/jvi.00712-11

60 Mänz, B. et al. Pandemic Influenza A Viruses Escape from Restriction by Human MxA through Adaptive Mutations in the Nucleoprotein. PLOS Pathogens 9, e1003279 (2013). 10.1371/journal.ppat.1003279

61 Deeg, C. M. et al. In vivo evasion of MxA by avian influenza viruses requires human signature in the viral nucleoprotein. J Exp Med 214, 1239–1248 (2017). 10.1084/jem.20161033

62 Haller, O. & Kochs, G. Mx genes: host determinants controlling influenza virus infection and trans-species transmission. Hum Genet 139, 695–705 (2020). 10.1007/s00439-019-02092-8

63 Cauldwell, A. V., Long, J. S., Moncorge, O. & Barclay, W. S. Viral determinants of influenza A virus host range. J Gen Virol 95, 1193–1210 (2014). 10.1099/vir.0.062836-0

64 Turnbull, M. L. et al. Role of the B Allele of Influenza A Virus Segment 8 in Setting Mammalian Host Range and Pathogenicity. J Virol 90, 9263–9284 (2016). 10.1128/JVI.01205-16

65 Nations, F. a. A. O. o. t. U. Livestock Systems, <https://www.fao.org/livestock-systems/global-distributions/cattle/en/> (2024).

66 Bourret, V. et al. Whole-genome, deep pyrosequencing analysis of a duck influenza A virus evolution in swine cells. Infection, Genetics and Evolution 18, 31–41 (2013). 10.1016/j.meegid.2013.04.034

67 Hardy, A. et al. The Timing and Magnitude of the Type I Interferon Response Are Correlated with Disease Tolerance in Arbovirus Infection. mBio 14, e0010123 (2023). 10.1128/mbio.00101-23

68 Yamada, S. et al. Biological and Structural Characterization of a Host-Adapting Amino Acid in Influenza Virus. PLOS Pathogens 6, e1001034 (2010). 10.1371/journal.ppat.1001034

69 Steel, J., Lowen, A. C., Mubareka, S. & Palese, P. Transmission of Influenza Virus in a Mammalian Host Is Increased by PB2 Amino Acids 627K or 627E/701N. PLOS Pathogens 5, e1000252 (2009). 10.1371/journal.ppat.1000252

70 Zhou, B. et al. Asparagine Substitution at PB2 Residue 701 Enhances the Replication, Pathogenicity, and Transmission of the 2009 Pandemic H1N1 Influenza A Virus. PLOS ONE 8, e67616 (2013). 10.1371/journal.pone.0067616

71 Zhang, X. et al. Enhanced pathogenicity and neurotropism of mouse-adapted H10N7 influenza virus are mediated by novel PB2 and NA mutations. Journal of General Virology 98, 1185–1195 (2017). 10.1099/jgv.0.000770

72 Gu, C. et al. A human isolate of bovine H5N1 is transmissible and lethal in animal models. Nature (2024). 10.1038/s41586-024-08254-7

73 Agüero, M. et al. Highly pathogenic avian influenza A(H5N1) virus infection in farmed minks, Spain, October 2022. Euro Surveill 28 (2023). 10.2807/1560-7917.Es.2023.28.3.2300001

74 Bouvier, N. M. & Lowen, A. C. Animal Models for Influenza Virus Pathogenesis and Transmission. Viruses 2, 1530–1563 (2010). 10.3390/v20801530

75 Hatta, M., Gao, P., Halfmann, P. & Kawaoka, Y. Molecular basis for high virulence of Hong Kong H5N1 influenza A viruses. Science 293, 1840–1842 (2001). 10.1126/science.1062882

76 Schrauwen, E. J. et al. The multibasic cleavage site in H5N1 virus is critical for systemic spread along the olfactory and hematogenous routes in ferrets. J Virol 86, 3975–3984 (2012). 10.1128/jvi.06828-11

77 Tipih, T. et al. Highly pathogenic avian influenza H5N1 clade 2.3.4.4b genotype B3.13 is highly virulent for mice, rapidly causing acute pulmonary and neurologic disease. Nat Commun 16, 5738 (2025). 10.1038/s41467-025-60407-y

78 Fabrizio, T. P. et al. Genotype B3.13 influenza A(H5N1) viruses isolated from dairy cattle demonstrate high virulence in laboratory models, but retain avian virus-like properties. Nature Communications 16, 6771 (2025). 10.1038/s41467-025-61757-3

79 Mostafa, A. et al. Avian influenza A (H5N1) virus in dairy cattle: origin, evolution, and cross-species transmission. mBio 15, e0254224 (2024). 10.1128/mbio.02542-24

80 Peacock, T. P. et al. The global H5N1 influenza panzootic in mammals. Nature 637, 304–313 (2025). 10.1038/s41586-024-08054-z

81 Halwe, N. J. et al. H5N1 clade 2.3.4.4b dynamics in experimentally infected calves and cows. Nature (2024). 10.1038/s41586-024-08063-y

82 Schoggins, J. W. Interferon-Stimulated Genes: What Do They All Do? Annu Rev Virol 6, 567–584 (2019). 10.1146/annurev-virology-092818-015756

83 Schoggins, J. W. et al. A diverse range of gene products are effectors of the type I interferon antiviral response. Nature 472, 481–485 (2011). 10.1038/nature09907

84 Shaw, A. E. et al. Fundamental properties of the mammalian innate immune system revealed by multispecies comparison of type I interferon responses. PLOS Biology 15, e2004086 (2017). 10.1371/journal.pbio.2004086

85 Ankerhold, J., Kessler, S., Beer, M., Schwemmle, M. & Ciminski, K. Replication Restriction of Influenza A (H5N1) Clade 2.3.4.4b Viruses by Human Immune Factor, 2023-2024. Emerg Infect Dis 31, 199-202 (2025). 10.3201/eid3101.241236

86 Briggs, K., Chrzastek, K., Segovia, K., Mo, J. & Kapczynski, D. R. Genetic insertion of mouse Myxovirus-resistance gene 1 increases innate resistance against both high and low pathogenic avian influenza virus by significantly decreasing replication in chicken DF1 cell line. Virology 595, 110066 (2024). 10.1016/j.virol.2024.110066

87 Wang, Y. et al. Associations of chicken Mx1 polymorphism with antiviral responses in avian influenza virus infected embryos and broilers. Poultry Science 91, 3019–3024 (2012). 10.3382/ps.2012-02471

88 Schusser, B. et al. Mx Is Dispensable for Interferon-Mediated Resistance of Chicken Cells against Influenza A Virus. Journal of Virology 85, 8307–8315 (2011). doi:10.1128/jvi.00535-11

89 Arnheiter, H. & Haller, O. Antiviral state against influenza virus neutralized by microinjection of antibodies to interferon-induced Mx proteins. Embo j 7, 1315–1320 (1988). 10.1002/j.1460-2075.1988.tb02946.x

90 Dornfeld, D., Petric Philipp, P., Hassan, E., Zell, R. & Schwemmle, M. Eurasian Avian-Like Swine Influenza A Viruses Escape Human MxA Restriction through Distinct Mutations in Their Nucleoprotein. Journal of Virology 93, 10.1128/jvi.00997-00918 (2019). 10.1128/jvi.00997-18

91 Riegger, D. et al. The nucleoprotein of newly emerged H7N9 influenza A virus harbors a unique motif conferring resistance to antiviral human MxA. J Virol 89, 2241–2252 (2015). 10.1128/jvi.02406-14

92 Ashenberg, O., Padmakumar, J., Doud, M. B. & Bloom, J. D. Deep mutational scanning identifies sites in influenza nucleoprotein that affect viral inhibition by MxA. PLoS Pathog 13, e1006288 (2017). 10.1371/journal.ppat.1006288

93 Fatima, U. et al. Equine Mx1 Restricts Influenza A Virus Replication by Targeting at Distinct Site of its Nucleoprotein. Viruses 11 (2019). 10.3390/v11121114

94 Gack, M. U. et al. Influenza A virus NS1 targets the ubiquitin ligase TRIM25 to evade recognition by the host viral RNA sensor RIG-I. Cell Host Microbe 5, 439–449 (2009). 10.1016/j.chom.2009.04.006

95 Hale, B. G., Randall, R. E., Ortin, J. & Jackson, D. The multifunctional NS1 protein of influenza A viruses. J Gen Virol 89, 2359–2376 (2008). 10.1099/vir.0.2008/004606-0

96 Pensaert, M., Ottis, K., Vandeputte, J., Kaplan, M. M. & Bachmann, P. A. Evidence for the natural transmission of influenza A virus from wild ducks to swine and its potential importance for man. Bull World Health Organ 59, 75–78 (1981).

97 Waddell, G. H., Teigland, M. B. & Sigel, M. M. A NEW INFLUENZA VIRUS ASSOCIATED WITH EQUINE RESPIRATORY DISEASE. J Am Vet Med Assoc 143, 587–590 (1963).

98 Webster, R. G. Are equine 1 influenza viruses still present in horses? Equine Vet J 25, 537–538 (1993). 10.1111/j.2042-3306.1993.tb03009.x

99 Guo, Y. et al. Characterization of a new avian-like influenza A virus from horses in China. Virology 188, 245–255 (1992).

100 Zhu, H., Hughes, J. & Murcia, P. R. Origins and Evolutionary Dynamics of H3N2 Canine Influenza Virus. Journal of virology 89, 5406–5418 (2015). 10.1128/jvi.03395-14

101 Plaza, P., Gamarra-Toledo, V., Euguí, J. R. & Lambertucci, S. Recent Changes in Patterns of Mammal Infection with Highly Pathogenic Avian Influenza A(H5N1) Virus Worldwide. Emerging Infectious Disease journal 30, 444 (2024). 10.3201/eid3003.231098

102 Uhart, M. M. et al. Epidemiological data of an influenza A/H5N1 outbreak in elephant seals in Argentina indicates mammal-to-mammal transmission. Nature Communications 15, 9516 (2024). 10.1038/s41467-024-53766-5

103 (!!! INVALID CITATION !!! 85).

104 Kareinen, L. et al. Highly pathogenic avian influenza A(H5N1) virus infections on fur farms connected to mass mortalities of black-headed gulls, Finland, July to October 2023. Eurosurveillance 29, 2400063 (2024). 10.2807/1560-7917.ES.2024.29.25.2400063

105 Mahase, E. H5N1: UK reports world’s first case in a sheep. Bmj 388, r591 (2025). 10.1136/bmj.r591

106 Yen, H. L. et al. The R292K mutation that confers resistance to neuraminidase inhibitors leads to competitive fitness loss of A/Shanghai/1/2013 (H7N9) influenza virus in ferrets. J Infect Dis 210, 1900–1908 (2014). 10.1093/infdis/jiu353

107 Subbarao, E. K., London, W. & Murphy, B. R. A single amino acid in the PB2 gene of influenza A virus is a determinant of host range. J Virol 67, 1761–1764 (1993). 10.1128/jvi.67.4.1761-1764.1993

108 Gilbertson, B. & Subbarao, K. Mammalian infections with highly pathogenic avian influenza viruses renew concerns of pandemic potential. Journal of Experimental Medicine 220(8):e20230447 (2023). 10.1084/jem.20230447

109 de Wit, E. et al. Efficient generation and growth of influenza virus A/PR/8/34 from eight cDNA fragments. Virus Res 103, 155–161 (2004). 10.1016/j.virusres.2004.02.028

110 Beare, A. S., Schild, G. C. & Craig, J. W. Trials in man with live recombinants made from A/PR/8/34 (H0 N1) and wild H3 N2 influenza viruses. Lancet 2, 729–732 (1975). 10.1016/s0140-6736(75)90720-5

111 Beare, A. S. & Hall, T. S. RECOMBINANT INFLUENZA-A VIRUSES AS LIVE VACCINES FOR MAN: Report to the Medical Research Council’s Committee on Influenza and other Respiratory Virus Vaccines. The Lancet 298, 1271–1273 (1971). 10.1016/S0140-6736(71)90597-6

112 Liu, M. et al. Human-type sialic acid receptors contribute to avian influenza A virus binding and entry by hetero-multivalent interactions. Nature Communications 13, 4054 (2022). 10.1038/s41467-022-31840-0

113 Steinegger, M. & Söding, J. MMseqs2 enables sensitive protein sequence searching for the analysis of massive data sets. Nat Biotechnol 35, 1026–1028 (2017). 10.1038/nbt.3988

114 Katoh, K. & Standley, D. M. MAFFT Multiple Sequence Alignment Software Version 7: Improvements in Performance and Usability. Molecular Biology and Evolution 30, 772–780 (2013). 10.1093/molbev/mst010

115 Minh, B. Q. et al. IQ-TREE 2: New Models and Efficient Methods for Phylogenetic Inference in the Genomic Era. Mol Biol Evol 37, 1530–1534 (2020). 10.1093/molbev/msaa015

116 Yu, G., Smith, D. K., Zhu, H., Guan, Y. & Lam, T. T.-Y. ggtree: an r package for visualization and annotation of phylogenetic trees with their covariates and other associated data. Methods in Ecology and Evolution 8, 28–36 (2017). 10.1111/2041-210X.12628

117 Hardy, A. et al. The Timing and Magnitude of the Type I Interferon Response Are Correlated with Disease Tolerance in Arbovirus Infection. mBio 14, e00101–00123 (2023). doi:10.1128/mbio.00101-23

118 Smith, M. C., Goddard, E. T., Perusina Lanfranca, M. & Davido, D. J. hTERT Extends the Life of Human Fibroblasts without Compromising Type I Interferon Signaling. PLOS ONE 8, e58233 (2013). 10.1371/journal.pone.0058233

119 Rihn, S. J. et al. A plasmid DNA-launched SARS-CoV-2 reverse genetics system and coronavirus toolkit for COVID-19 research. PLOS Biology 19, e3001091 (2021). 10.1371/journal.pbio.3001091

120 Hoffmann, E., Stech, J., Guan, Y., Webster, R. G. & Perez, D. R. Universal primer set for the full-length amplification of all influenza A viruses. Arch Virol 146, 2275–2289 (2001).

121 Bakshi, S., Taylor, J., Strickson, S., McCartney, T. & Cohen, P. Identification of TBK1 complexes required for the phosphorylation of IRF3 and the production of interferon β. Biochem J 474, 1163–1174 (2017). 10.1042/bcj20160992

122 Varela, M. et al. Schmallenberg Virus Pathogenesis, Tropism and Interaction with the Innate Immune System of the Host. PLOS Pathogens 9, e1003133 (2013). 10.1371/journal.ppat.1003133

123 Meehan, G. R. et al. Phenotyping the virulence of SARS-CoV-2 variants in hamsters by digital pathology and machine learning. PLOS Pathogens 19, e1011589 (2023). 10.1371/journal.ppat.1011589

124 Bankhead, P. et al. QuPath: Open source software for digital pathology image analysis. Scientific Reports 7, 16878 (2017). 10.1038/s41598-017-17204-5

125 Zhang, X. et al. Enhanced pathogenicity and neurotropism of mouse-adapted H10N7 influenza virus are mediated by novel PB2 and NA mutations. J Gen Virol 98, 1185–1195 (2017). 10.1099/jgv.0.000770

126 Bussey, K. A., Bousse, T. L., Desmet, E. A., Kim, B. & Takimoto, T. PB2 residue 271 plays a key role in enhanced polymerase activity of influenza A viruses in mammalian host cells. J Virol 84, 4395–4406 (2010). 10.1128/jvi.02642-09

127 Zhou, B. et al. PB2 Residue 158 Is a Pathogenic Determinant of Pandemic H1N1 and H5 Influenza A Viruses in Mice. Journal of Virology 85, 357–365 (2011). 10.1128/jvi.01694-10

128 Fan, S. et al. Novel residues in avian influenza virus PB2 protein affect virulence in mammalian hosts. Nature Communications 5, 5021 (2014). 10.1038/ncomms6021

129 Peacock, T. P. et al. Mammalian ANP32A and ANP32B Proteins Drive Differential Polymerase Adaptations in Avian Influenza Virus. Journal of Virology 97, e00213–00223 (2023). doi:10.1128/jvi.00213-23

130 Xiao, C. et al. PB2-588 V promotes the mammalian adaptation of H10N8, H7N9 and H9N2 avian influenza viruses. Scientific Reports 6, 19474 (2016). 10.1038/srep19474

